# Seasonal shifts in the gut microbiome indicate plastic responses to diet in wild geladas

**DOI:** 10.1101/2020.07.07.192336

**Authors:** Alice Baniel, Katherine R Amato, Jacinta C Beehner, Thore J Bergman, Arianne Mercer, Rachel F Perlman, Lauren Petrullo, Laurie Reitsema, Sierra Sams, Amy Lu, Noah Snyder-Mackler

**Affiliations:** Department of Anthropology, Stony Brook University, Stony Brook, NY 11794, USA; Department of Anthropology, Northwestern University, Evanston, IL 60208, USA; Department of Psychology, University of Michigan, Ann Arbor, MI 48109, USA; Department of Anthropology, University of Michigan, Ann Arbor, MI 48109, USA; Department of Ecology and Evolutionary Biology, University of Michigan, Ann Arbor, MI 48109, USA; Department of Psychology, University of Washington, Seattle, WA 98195, USA; Interdepartmental Doctoral Program in Anthropological Sciences, Stony Brook University, Stony Brook, NY 11794, USA; Department of Anthropology, University of Georgia, Athens, GA 30602, USA; Center for Evolution and Medicine, Arizona State University, Tempe, AZ 85281, USA; School of Life Sciences, Arizona State University, Tempe, AZ 85287, USA; Department of Biology, University of Washington, Seattle, WA 98195, USA

**Keywords:** gut microbiome, graminivory, seasonality, thermoregulation, Theropithecus geladas, primates

## Abstract

Animals have evolved numerous strategies to cope with energetic challenges, with dynamic changes to the gut microbiome potentially constituting one such strategy. We tested how proxies of food availability (rainfall) and thermoregulatory stress (temperature) predicted gut microbiome composition of geladas (*Theropithecus geladas*), a grazing, high-altitude primate inhabiting a seasonal environment. The gelada gut microbiome varied across seasons, reflecting more efficient digestion of the primary foods eaten at certain times of year. In rainier periods, the gut was dominated by cellulolytic/fermentative bacteria that specialized in digesting grass, while during dry periods the gut was dominated by bacteria that break down starches found in underground plant parts. Temperature had a smaller, but detectable, effect on the gut microbiome. We found an increase in microbes involved in metabolism and energy production during cold and dry periods, suggesting buffering when thermoregulatory and nutritional stress co-occurred. Our results suggest that the gelada gut microbiome may shift to compensate for host diet and energetic demands.

## INTRODUCTION

Obtaining sufficient nutrients is a fundamental challenge for most animals. Yet, the availability and nutritional content of food can vary temporally and spatially in response to changes in climate and geography. Nutritional demands further vary in response to thermoregulatory needs and life history processes, such as growth and reproduction (McNab 2002; Dufour and Sauther 2002). Animals have evolved a variety of behavioral and physiological strategies to cope with these shifting demands, including altered feeding and activity patterns and increased mobilization of stored fat to fuel energetic demands (Doran 1997; Gursky 2000; van Schaik and Brockman 2005; Dias et al. 2011). Recently, the gut microbiome has been proposed as an additional avenue by which animals can cope with changing dietary landscapes and energetic challenges (Candela et al. 2012; David et al. 2014; Amato et al. 2015). The gastrointestinal tract of animals harbors a dense microbial community that helps to break down and ferment plant structural carbohydrates, producing short-chain fatty acids (SCFAs) that can be used as an energy source by hosts (Bäckhed 2011; Flint et al. 2012; White et al. 2014). The absorption of SCFAs in the gut may be particularly important for herbivorous species, such as foregut and hindgut fermenters, which obtain as much as 40-90% of their energy requirements from bacterial degradation of complex plant polysaccharides (Bergman et al. 1965; Udén et al. 1982; Milton and McBee 1983; Popovich et al. 1997). Additionally, variation in gut microbiome composition affects the efficiency of caloric harvest and the metabolic programming of the host (De Filippo et al. 2010; Bäckhed 2011; Krajmalnik-Brown et al. 2012; Tremaroli and Bäckhed 2012; Hanning and Diaz-Sanchez 2015). For instance, in mice (*Mus musculus*) and humans, obese and lean individuals have strikingly different gut microbiota composition, with obese phenotypes being associated with higher energy extraction from diet and increased lipogenesis (Turnbaugh et al. 2006; Turnbaugh and Gordon 2009; Tseng and Wu 2019).

In wild mammals, the gut microbiome responds rapidly to seasonal and dietary changes (Maurice et al. 2015; Liu et al. 2019; Sun et al. 2016; Amato et al. 2015; Mallott et al. 2018; Ren et al. 2016; Springer et al. 2017), presumably to buffer seasonal energetic challenges (Amato et al. 2015; Sun et al. 2016). For example, a simultaneous increase in bacterial taxa involved in fiber fermentation and in SCFA concentrations during the dry season were suggested to allow Mexican black howler monkeys (*Alouatta pigra*) to maintain energy balance during energetic shortfalls without changes in activity or ranging patterns (Amato et al. 2015). Moreover, gut bacteria increase intestinal absorptive capacity, energy homeostasis, and fat burning during cold periods in mice (Chevalier et al. 2015), and improve digestive efficiency and SCFA production in energetically challenged ruminants living at cold and high-altitude (Zhang et al. 2016; Li et al. 2018). These microbial shifts likely come at some cost. For instance, increases in microbes that improve host metabolism under certain conditions may reduce the abundance of microbes that support host immune function (Amato et al. 2014; Reese and Kearney 2019). However, in seasonal and nutritionally challenging environments, enduring these trade-offs may be necessary for host survival and reproduction.

Geladas (*Theropithecus gelada*) represent an excellent system to investigate the relationship between gut microbiota composition and seasonal variation in host diet and energy needs. Despite being the only graminivorous primate with up to 90% of their diet comprised of grass (Fashing et al. 2014; Jarvey et al. 2018), their gastrointestinal tract appears poorly adapted to this specialization (but see (Wrangham 1980; Venkataraman et al. 2014) for dental, manual, and locomotor adaptations), closely resembling their closest phylogenetic relatives, baboons (*Papio* spp.) – a taxon that is omnivorous (Mau et al. 2011). To compensate, geladas may rely heavily on their gut microbiota to maximize nutrient extraction from grasses, likely through hindgut fermentation (Mau et al. 2011; Trosvik et al. 2018). Moreover, geladas live in a high-altitude, energetically demanding environment that exhibits marked inter- and intra-annual fluctuation in rainfall and temperature (Jarvey et al. 2018; Tinsley Johnson et al. 2018). During rainier months, when grass is abundant, they focus almost exclusively on eating above-ground graminoid leaves and seeds, and during drier months, when grass availability decreases, they shift heavily to underground foods (rhizomes, roots, corms, bulbs) (Hunter 2001; Jarvey et al. 2018). This diet provides distinct challenges. Underground foods are considered a fallback food for geladas since they take additional time and effort to harvest, are harder to process, and are relied upon only when grasses are less abundant (Venkataraman et al. 2014; Jarvey et al. 2018). Despite being considered a fallback food, these underground foods are rich in starches and carbohydrates, suggesting that they contain more nutritional energy than grass (Dominy et al. 2008). This high amount of energy, however, comes at some cost: roots and rhizomes are generally higher in fibers and lignin - and thus harder to digest than grasses. In addition to these nutritional challenges, ambient temperatures frequently drop to near freezing in some months, and the metabolic costs of thermoregulation are known to strongly influence gelada physiology (Beehner and McCann 2008) and the timing of reproduction (Tinsley Johnson et al. 2018; Carrera et al. 2020). Thus, seasonal dietary shifts and temperature variation may entail distinct digestive and thermoregulatory challenges.

One previous study on geladas from Guassa, Ethiopia found that gut microbial communities of adult females do indeed shift across seasons (Trosvik et al. 2018), supporting the hypothesis that the gut microbiome may help hosts confront environmental challenges. This study focused on adult females and assessed seasonal variation by separating the samples into two categorical seasons (i.e., rainy, dry). Our study expands on this study by including adult males, incorporating continuous climatic data across several years, and examining proxies of thermoregulatory stress (in addition to diet) as factors that can influence the composition and function of the gelada gut microbiome. Indeed, rainfall and temperature vary independently of each other and represent distinct ecological challenges in gelada ecosystems. Therefore, we were interested in further testing which aspect of gelada ecology more strongly determines seasonal microbiome shifts.

We analysed the gut microbiome composition and predicted microbiome function in 758 fecal samples across 5 years from 48 adult male and 86 adult female geladas living in the Ethiopian highlands in the Simien Mountains National Park. The Simien Mountains Gelada Research Project (SMGRP) has been collecting detailed climatologic, demographic, and behavioral data from this study population since 2006, allowing us to examine how ecological (rainfall and temperature) and individual (group membership, sex, reproductive status, and age) factors influence gelada gut microbiome composition. We hypothesized that ecological factors would be more strongly associated with variation in the gelada microbiome than individual factors, and that rainfall and temperature would have independent effects. In particular, we expected that rainfall, which is a good proxy for grass availability (Jarvey et al. 2018), would have the strongest effect on the gelada gut microbiome. Specifically, we predicted that the taxonomic changes associated with rainfall would mainly reflect a shift to grass-based versus underground food-based diet, in order to allow individuals to maximize energy extraction from those seasonal foods. We found that the gelada microbiome exhibited drastic shifts related to climatological variables; but individual variables, like age and sex, had minimal effects. Rainfall and temperature exerted independent effects on the microbial composition and predicted function – with rainfall having a stronger effect on the gelada gut bacteria. High rainfall, which is correlated with grass availability (Jarvey et al. 2018), was associated with more cellulolytic and fibrolytic bacterial taxa, when graminoid leaves were the main food source. Dry periods, which are correlated to underground food consumption (Jarvey et al. 2018), were associated with amylolytic and methanogenic taxa. Cold periods were further characterized by more amylolytic taxa, and hot periods by more methanogenic taxa. In both drier and colder periods, the gut microbiome shifted to predicted functions that suggested increased digestive efficiency, including energy, amino acid and lipid metabolism. Overall, gelada gut microbial composition covaried with diet and temperature in a pattern that suggests plastic but distinct responses to different dietary and metabolic challenges.

## RESULTS

### The gelada gut microbiome

We identified 3,295 amplicon specific variants (ASVs) in 758 fecal samples (mean±SD=813±243 ASVs per sample, range=92-1730) using deep 16S rRNA gene amplicon sequencing. These 3,295 ASVs came from 16 different phyla, 65 families, and 200 genera (Table S1, Figure 1, Figures S1-S2). Of the 3,295 ASVs, 170 (5%) were present in at least 90% of samples and form what can be considered the “core microbiota” of geladas (Table S2). The four most abundant bacterial phyla were *Firmicutes* (32%), *Kiritimatiellaeota* (formerly called *Verrucomicrobiota subdivision 5;* 26%), *Bacteroidetes* (23%), and *Spirochaetes* (5%) (Table S1, Figure 1A). All microbes assigned to *Kiritimatiellaeota* were part of the *RFP12* family and represent almost one quarter of the gelada gut microbiome (mean 26%, range 0.02%-70%, Figure 1B). Although the metabolic function of *RFP12* remains unknown, those bacteria have been found in high quantities in the gut of some domestic horse (*Equus ferus caballus*) and sheep (*Ovis aries*) populations (Steelman et al. 2012; Costa et al. 2015; Wang et al. 2017) and could have some fermentative function. Other taxa found at high frequency in the guts of ruminants and herbivorous hindgut fermenters were also prevalent in the gelada gut, including many cellulolytic/fibrolytic (13% *Ruminococcaceae*, 6% *Lachnospiraceae*, 4% *Clostridiales vadinBB60 group*, 1.5% *Fibrobacteraceae*) and fermentative families (*5.3% Rikenellaceae*, 5% *Prevotellaceae, 4.1% Bacteroidales F082*) (Table S1, Figure 1B and S1). The *Spirochaetes* phylum was mostly composed of *Treponema* (3.5%), a genus involved in lignocellulose degradation (Warnecke et al. 2007).

**Figure 1.**
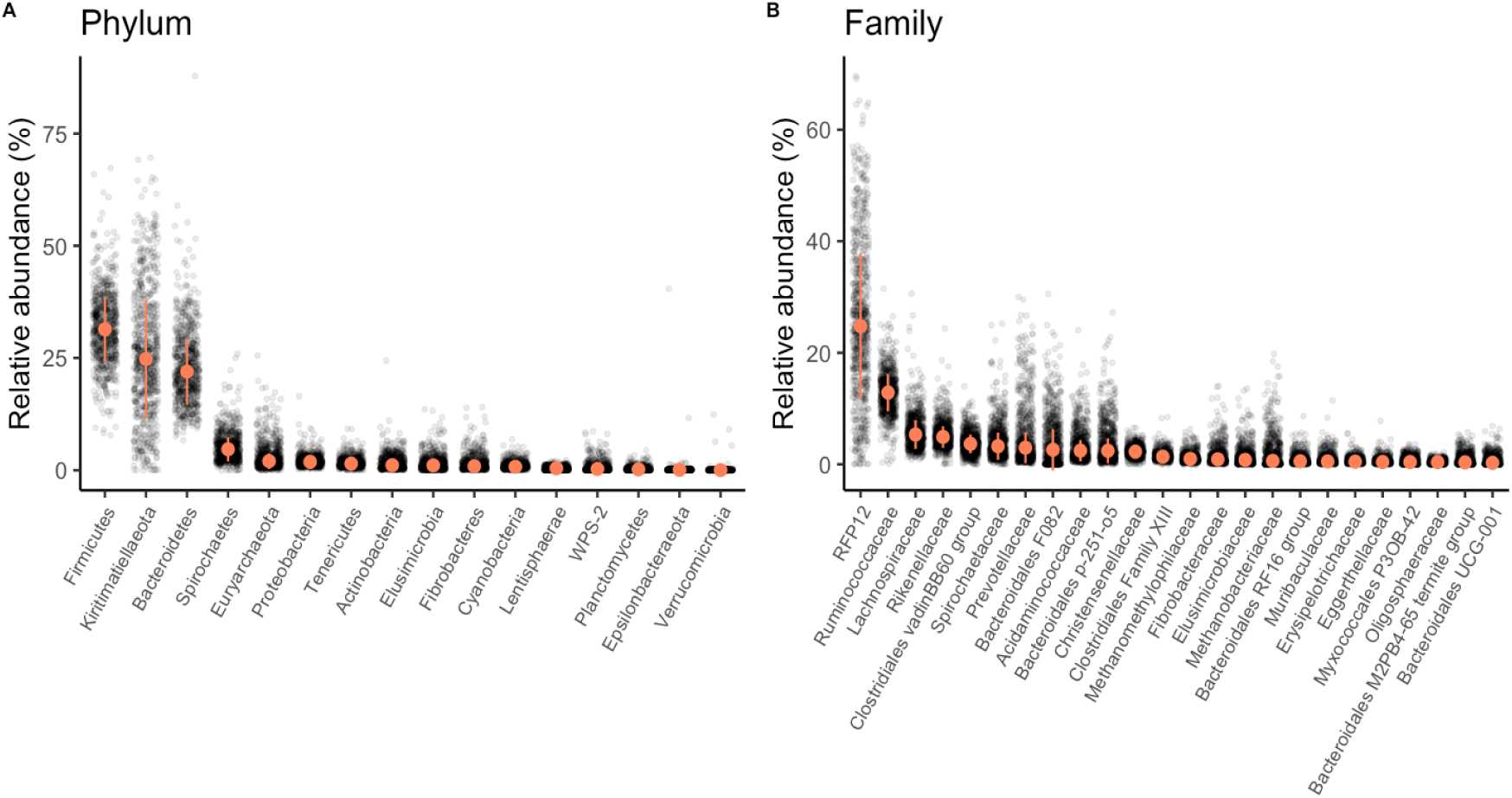
Taxonomic composition of the geladas gut at the phylum and family levels. Relative abundance **(A)** of all bacterial phyla and **(B)** of the 24 most abundant families (whose relative abundance>0.02%) in the gelada feces. The median and median absolute deviation (error limit) are represented in orange.

### Dietary changes

To examine how seasonal variation in rainfall and temperature was associated with changes in the gelada gut microbiome, we used measures of true climatic conditions, including monthly cumulative rainfall (an appropriate proxy of grass availability in the Simiens: (Jarvey et al. 2018)) and average monthly minimum temperature (a proxy of thermoregulatory constraint: (Beehner and McCann 2008; Tinsley Johnson et al. 2018)). At the level of within-sample community diversity (“alpha diversity”), we found that cumulative rainfall was positively associated with Shannon evenness (Table 1, Figure 2A,C) but had no effect on bacterial richness or Faith’s phylogenetic diversity (Table S3, Figure S3). Thus, rainfall was associated with the relative abundance of ASVs within a sample but not the absolute number of ASVs or their phylogenetic diversity.

**Table 1.** Determinants of alpha diversity, as measured by the Shannon index.

| Fixed factor | Estimate | SE | 95% confidence interval | LRT | P-value |
| --- | --- | --- | --- | --- | --- |
| **Sex (male)** | -0.12 | 0.04 | [-0.20 ; -0.05] | 9.27 | **0.002** |
| Age | 0.01 | 0.02 | [-0.02 ; 0.04] | 0.49 | 0.484 |
| **Cumulative rainfall** | 0.05 | 0.02 | [0.02 ; 0.08] | 12.13 | **<0.001** |
| Min temperature | 0.00 | 0.01 | [-0.03 ; 0.03] | 0.04 | 0.848 |
| **Sequencing depth** | 0.07 | 0.01 | [0.04 ; 0.10] | 24.24 | **<0.001** |

Parameters and tests are based on linear mixed models of 758 samples and 131 individuals, controlling for individual identity and unit membership. Factors with p-values less than 0.05 are highlighted in bold.

**Figure 2.**
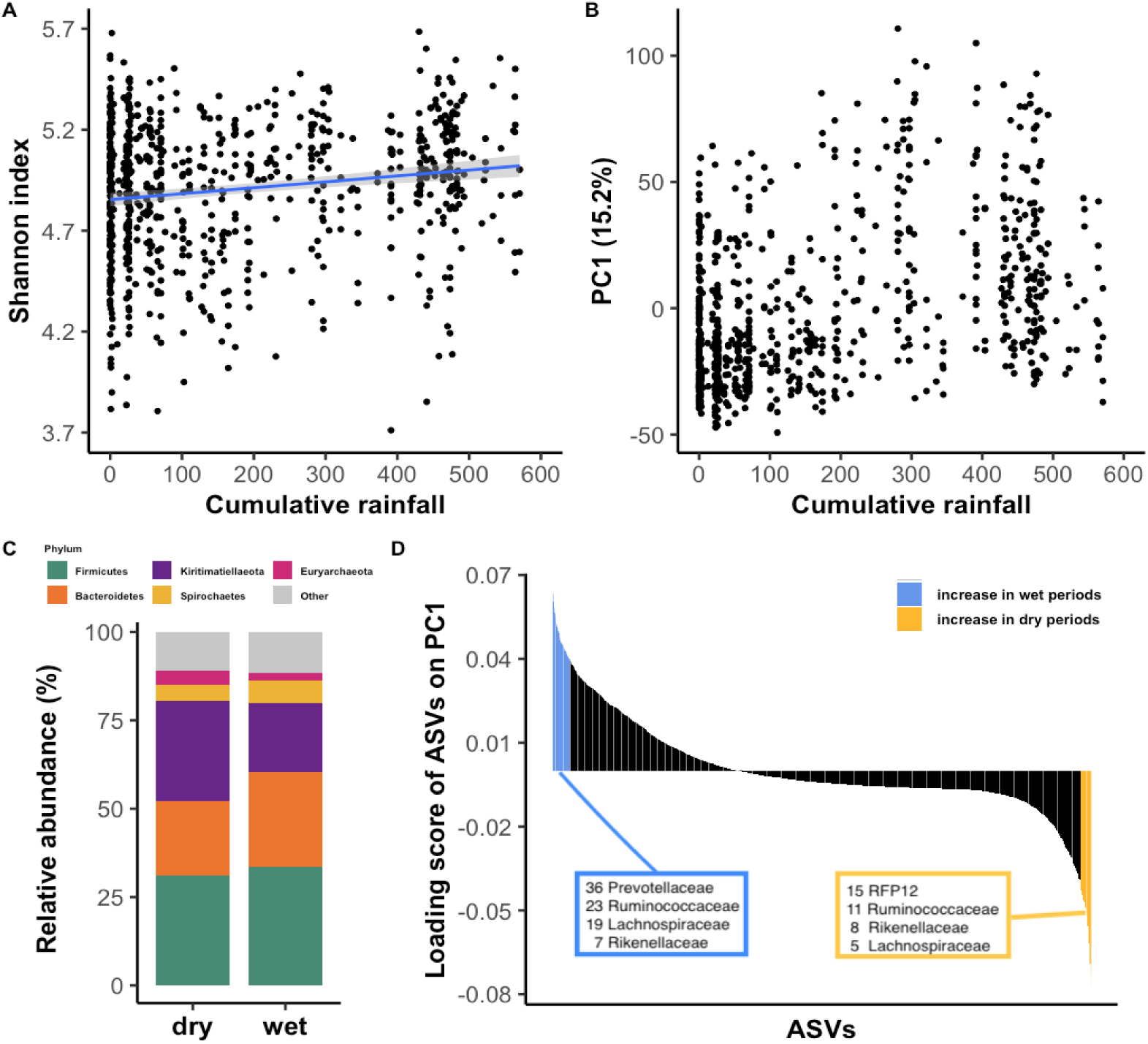
Rainfall structures the gelada gut microbiome. **(A)** Partial residual plot of Shannon alpha diversity index according to cumulative rainfall (in mm). Black dots represent the partial residuals from the LMM (i.e. showing the association between cumulative rainfall and alpha diversity, while controlling for all other predictors). The blue line and confidence intervals come from a linear regression (for representation only). Seven outlier samples (with a particularly low Shannon index) were omitted for clarity of representation. **(B)** Visualization of between-sample dissimilarity (based on Aitchison distance) on the first principal component (PC1) according to cumulative rainfall. **(C)** Compositional barplot of the five most abundant phyla in the dry (<100mm of rain in the past month, N=362) and wet (>200mm of rain in the past month, N=282) samples (cumulative rainfall was converted to a categorical variable for representation purposes). **(D)** Loading scores of each amplicon sequence variant (ASV) on the first principal component. ASVs with a loading score >0.4 (characteristics of the wet season) and <-0.4 (characteristic of the dry season) are colored.

Cumulative rainfall significantly explained 3.3% of the overall compositional dissimilarity - or beta diversity - between samples (as measured by Aitchison distance) (Table 2), which was nonetheless less than that explained by two demographic variables, individual identity and unit (social group) membership (20% and 6%, respectively; Table 2). The first principal component of beta diversity, which explained 15% of variation, was strongly associated with rainfall (r=0.43, t=12.93, df=756, p<0.001, Figure 2B). The ASVs that loaded positively on PC1 (i.e. correlated with higher rainfall, Figure 2D) were primarily from the families *Prevotellaceae, Ruminococcaceae*, and *Lachnospiraceae* (Table S4 and S5). By contrast, the ASVs that loaded negatively on PC1 (i.e. more abundant in low rainfall, Figure 2D) belonged to the family *RFP12* and a different subset of *Ruminococcaceae* that were not abundant during the wet season (Table S4 and S5).

**Table 2.** Results of PERMANOVA testing for the predictors that significantly structure the gut microbiome of geladas, using 10,000 permutations and the Aitchison dissimilarity distance between samples. The R-squared values indicate the amount of between-sample variation explained by each variable.

| Factor | R2 (%) | P-value |
| --- | --- | --- |
| **Individual<sup>1</sup>** | 20.25 | **<0.001** |
| **Sequencing depth<sup>2</sup>** | 3.77 | **<0.001** |
| **Unit<sup>2</sup>** | 5.84 | **<0.001** |
| **Cumulative rainfall<sup>2</sup>** | 3.30 | **<0.001** |
| **Min temperature<sup>2</sup>** | 0.33 | **<0.001** |
| **Sex<sup>2</sup>** | 0.23 | **0.012** |
| **Age<sup>2</sup>** | 0.19 | 0.045 |

<sup>1</sup> We first fit a model with individual identity as the only predictor in a PERMANOVA to estimate the sole effect of individual identity at explaining the overall gut composition of samples.
<sup>2</sup> We then fit a second PERMANOVA model where all other predictors were fit, stratifying on individual identity to control for pseudoreplication of samples from the same individual.

Cumulative rainfall predicted the relative abundance of gut microbes at all taxonomic levels and was significantly associated with the relative abundance of 63% of bacterial families tested (59-81% of taxa at other taxonomic levels, Figure 3, pBH<0.05). Thus, across most taxa, there was a clear contrast in the relative abundance of gut bacteria between the wet and dry periods (Table S6, Figure 4). In wetter periods, there was an increase in several important fermentative families from the *Bacteroides* order (including *Prevotellaceae* and *Bacteroidaceae)*, as well as in several cellulolytic/fibrolytic taxa (*Lachnospiraceae, Fibrobacteraceae, Spirochaetaceae* and several genera from the *Ruminococcaceae*; Figure 4 and 5A), suggesting improved digestive efficiency of plant cell wall polysaccharides at a time when the gelada diet consists mainly of grasses. In particular, nine *Prevotella* genera as well as the *Bacteroides* genus were higher during wetter periods than drier periods (Table S6, Figure 4B). There was also an increase in several proficient cellulolytic genera (e.g. *Senegalimassilia, Butyrivibrio, Saccharofermentans, Cellulosilyticum, Marvinbryantia*) (Table S6, Figure 4B). By contrast, the dry season was characterized by an increase in amylolytic genera (*Succinivibrio*; *Streptococcus* and *Pirellulaceae p-1088-a5 gut group*), in several efficient sugar-fermenting families (*Victivallales vadinBE97, Christensenellaceae*), and in the methane-producer *Methanobrevibacter*, a genus known to increase the rate of fermentation and digestive efficiency (Table S6, and Figures 4 and 5B). Consistent with our beta diversity analyses, we also found an increase in the relative abundance of the *RFP12* family during the dry season (Table S6, Figure 5B).

**Figure 3.**
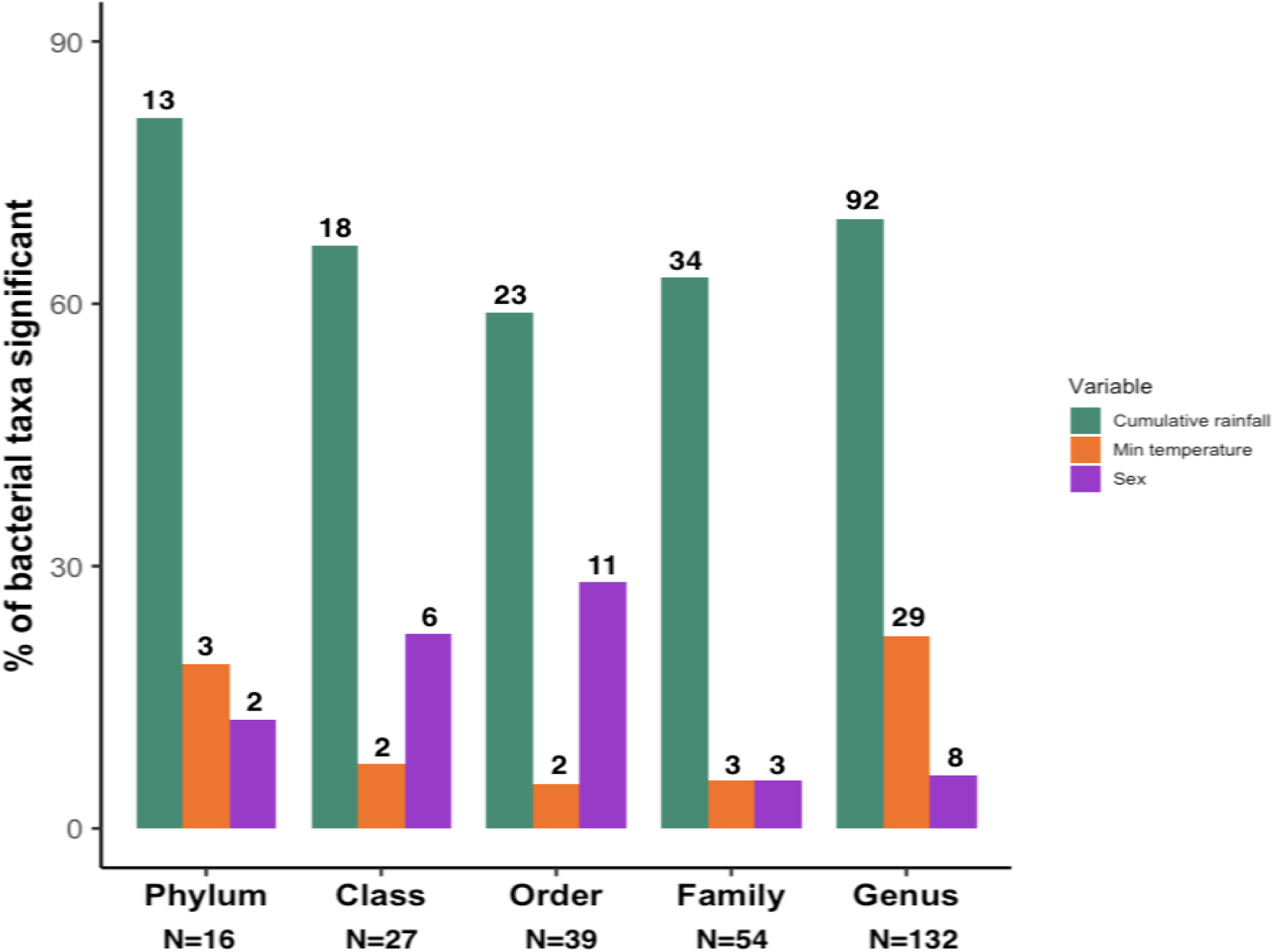
Rainfall exerts the strongest effect on bacterial relative abundance. Percent of taxa that are significantly associated (p_BH_<0.05) with rainfall (purple bars), temperature (orange bars), or sex (green bars), across five taxonomic levels. For a given bacterial taxa, the significance of each predictor was assessed using a negative binomial GLMM fitted on the count of this taxa per sample (controlling for sequencing depth as an offset factor, and including individual and unit membership as random effects). Only taxa with p_BH_ < 0.05 were considered significant. The numbers above the bars depict the number of taxa significantly differentially abundant, while the numbers below indicate the total taxa measured per level. Age was not significantly associated with relative abundance of any taxa at any level.

**Figure 4.**
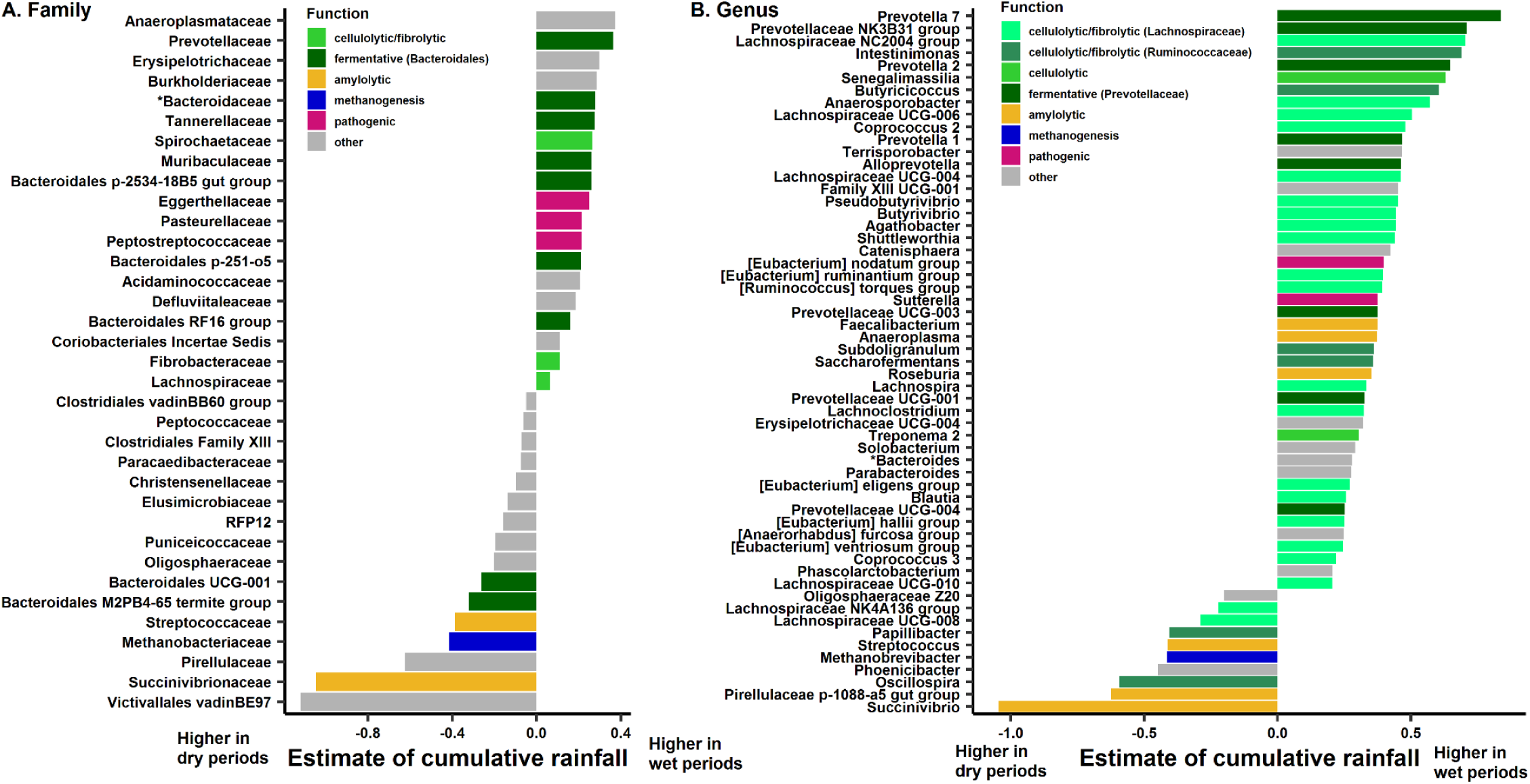
Rainfall predicts the relative abundance of many bacterial taxa. **(A)** Families and **(B)** Genera that are found differentially abundant according to cumulative rainfall. The estimate for each taxa of the cumulative rainfall effect comes from a negative binomial GLMM fitted on the count of this taxa per sample (controlling for sequencing depth as an offset factor, and including individual and unit membership as random effects). Taxa starting with “*” were fit with a binomial model instead. Only taxa with p_BH_ < 0.05 were considered significant. For ease of representation on panel B, only genera with effect sizes > |0.2| are represented. The full list of differentially abundant genera can be found in Table S8. Assignment of the “broad function” of a family or genus is for representation only, and is a simplification of the various functions subsumed within each taxonomic group.

**Figure 5.**
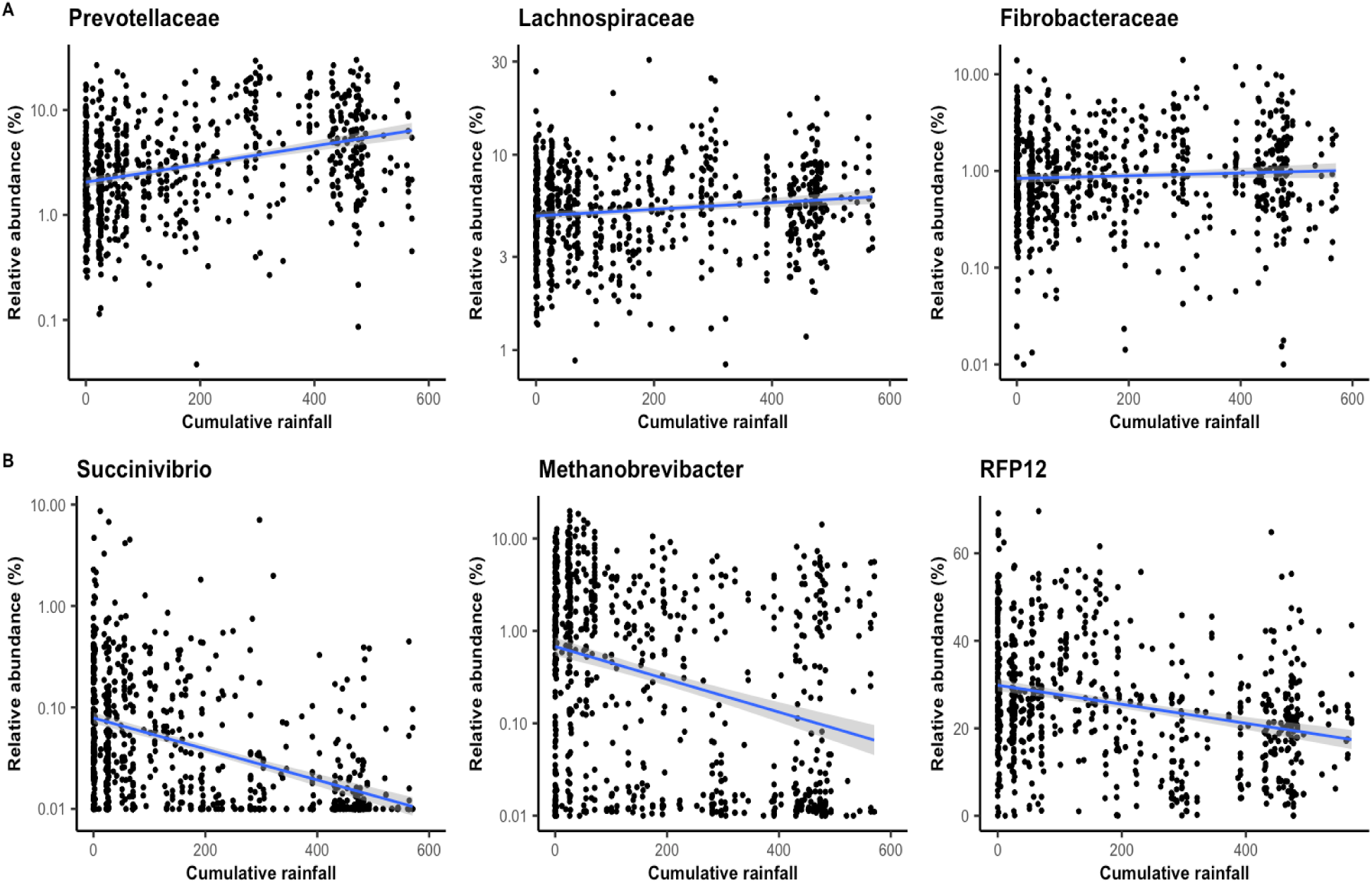
Relative abundance in six bacterial taxa (family or genus) that are significantly associated with rainfall. **(A)** more abundant during the wet season and **(B)** more abundant during the dry season. Note that the tick marks on the y-axis are spaced on a log10 scale (except for RFP12 which is plotted on a raw scale because of its high abundance). The blue line and confidence intervals come from a linear regression (for representation only). The significance of those effects have been estimated using negative binomial GLMMs including individual and unit membership as random effects.

The taxonomic changes associated with rainfall also corresponded to changes in the predicted function of the gelada gut microbiome (as assessed by PICRUSt2: (Douglas et al. 2019), NSTI mean±SD=0.60±0.13). During the wetter periods, functional changes tended to reflect the activity of the cellulolytic and fermentative bacterial taxa. Microbial pathways involved in the transport of molecules through bacterial membranes (e.g. ions, sugars, lipids, peptides), DNA replication and repair, and cell motility (Tables S7-S8, Figure 6, S4 and S5A) increased. We further found an increase in the metabolism of sugars (e.g. starch and sucrose metabolism, fructose, mannose, and galactose) (Figure S4 and S5A). Such activity probably reflects the exportation of sugar-cleaving enzymes and cellulosome complex across the outer membrane(Biddle et al. 2013; White et al. 2014) of fibrolytic bacteria (complex polysaccharides are too big to penetrate directly inside bacteria and have to be cleaved first) and the absorption of the soluble oligosaccharides back across the bacterial membrane (Biddle et al. 2013; White et al. 2014).

**Figure 6.**
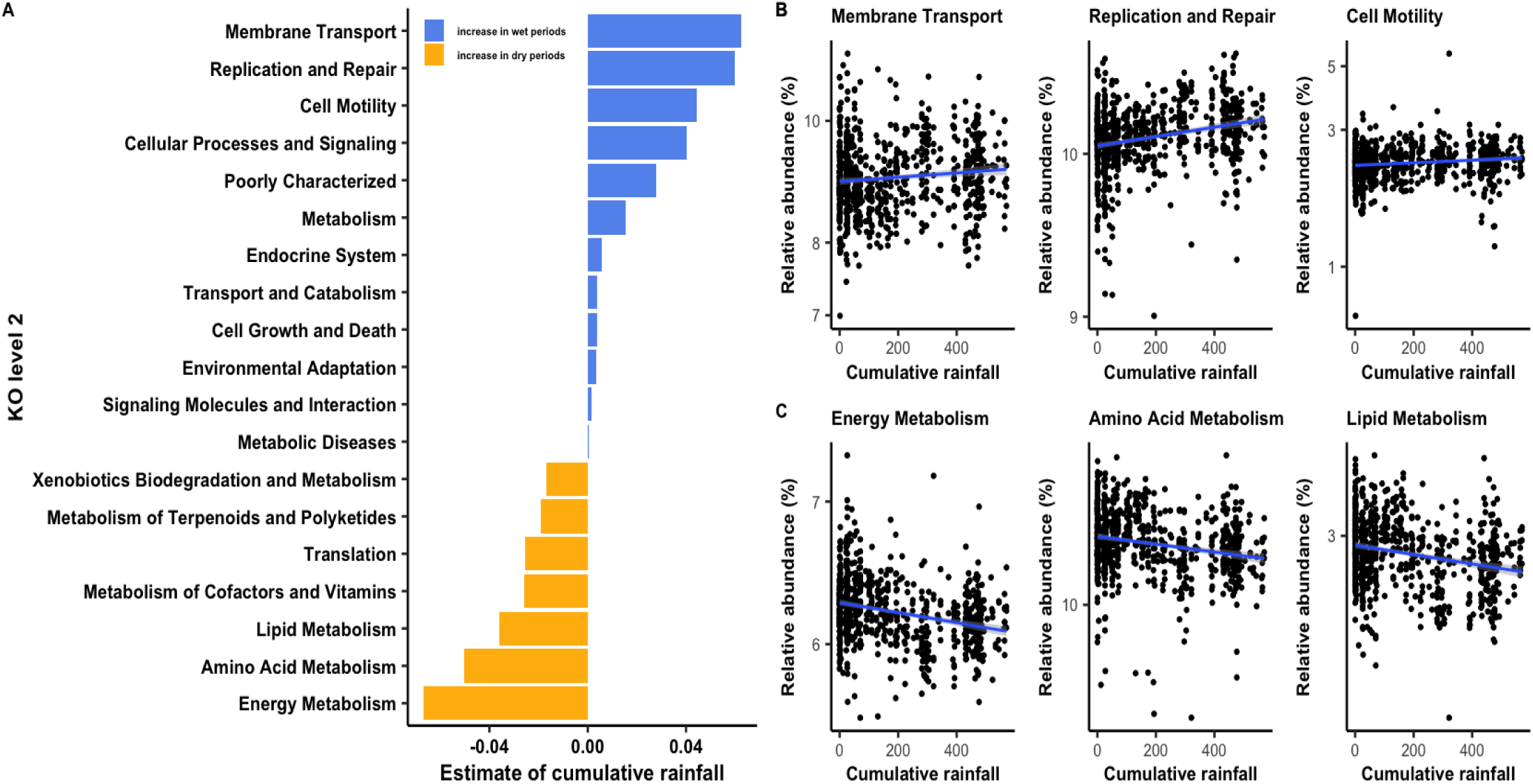
Rainfall predicts the functional profile of the gut microbiome. **(A)** Bacterial pathways at level 2 of KEGG Orthology (KO) that are differentially abundant according to cumulative rainfall (in mm). The estimate of the “rainfall” effect for each pathway comes from a LMM fitted on the relative abundance of each pathway per sample. Only pathways with pBH<0.05 are reported. Relative abundance of the three most enhanced functional pathways during **(B)** the wet season and **(C)** the dry season according to monthly cumulative rainfall. Note that the tick marks on the y-axis are spaced on a log10 scale. The blue line and confidence intervals come from a linear regression (for representation only). The significance of the rainfall effect effects per pathway have been estimated using LMMs including individual and unit membership as random effects.

During drier periods, the gelada gut harbored a greater abundance of bacterial genes involved in energy, amino acid, and lipid metabolism (Tables S7-S8 and Figure 6A,C). In particular, cellular energy production and cellular activity were enhanced during this period, as evidenced by increases in pathways involved in the citric acid cycle, oxidative phosphorylation, and fatty acid synthesis and metabolism (Figure S4 and S5B). Other energy metabolism pathways also increased during drier periods, including the methane pathway and the carbon fixation pathways, which are important for generating energy in anaerobic bacteria (Figure S4 and S5B). Finally, drier periods were associated with an increase in functions related to the synthesis of proteinogenic amino acids (e.g. tryptophan), the translation and synthesis of proteins (Figure S4), and the synthesis of lipopolysaccharide.

### Temperature

Compared to rainfall, minimum temperature had a much smaller impact on the gut microbiome. Average minimum temperature did not influence any metric of alpha diversity (Table 1 and S3, Figure S6A), and explained only 0.33% of the variation in beta diversity (Table 2, Figure S6B). Changes in temperature were significantly associated with the relative abundance of 5% of the families (5-22% at other taxonomic levels; Figure 3; pBH<0.05). More specifically, colder temperatures were characterized by a greater abundance of two amylolytic genera (*Lactobacillus* and *Streptococcus*); in several sugar-fermenting (*Hydrogenoanaerobacterium, Clostridium sensu stricto 1, Coprococcus 1*) and cellulose-degrading bacteria (*Marvinbryantia* and two genera from the *Ruminococcaceae* family) (Table S6, Figure S7). By contrast, hotter temperatures were associated with an increase in *Verrucomicrobia*, in the methane-producer *Methanobrevibacter*, and in several cellulolytic/fibrolytic genera from the *Ruminococcaceae and Lachnospiraceae* families (Table S6, Figure S7).

Similar to our taxonomic analysis, we found that temperature had a much smaller effect of the predicted function of the gelada gut microbiome (Tables S7-S8, Figure S8). During colder periods, we found a predicted increase in bacterial pathways involved in lipid metabolism and energy production (notably in oxidative phosphorylation pathway; Figure S8). Other pathways that increased during colder periods involved DNA repair and recombination and the bacterial secretion system. During hotter weather, pathways were more-poorly characterized and less specific, with predicted increases in methane metabolism and ABC transport (a membrane transporter).

### Sex, reproductive state, and age

The gut microbiome of females exhibited higher alpha diversity compared to males, regardless of the metric (richness, evenness, and Shannon index) (Table 1 and S3, Figure S9A). Across samples however, sex explained little between-sample variation (i.e. <1%) (Table 2, Figure S9B). We detected a handful of bacterial taxa that were differentially abundant according to sex (Table S6, Figure 3). At the phylum level, females harboured more *Verrucomicrobia* and *Proteobacteria* (particularly from class *Gammaproteobacteria, Deltaproteobacteria* and *Alphaproteobacteria*). At the family and genus levels, females had more taxa involved in lactic acid metabolism (*Lactobacillaceae, Anaerovibrio)*, cellulolysis (*Saccharofermentans*) and regulation of glucose and fat transpor*t* (*Erysipelatoclostridium*). Males, on the other hand, only harboured more *Pirellulales*. No predicted metabolic pathway was found to be differentially abundant with sex (Tables S7-S8).

Female reproductive state did not influence any alpha diversity metric (Table S9, Figure S10A) and was not a significant factor influencing beta diversity between samples (Table S10, Figure S10B). Very few taxa were differentially abundant according to female reproductive state (Table S11 and S12). Pregnant females harboured more *Verrucomicrobiota* (class *Verrucomicrobiae*) and *Epsilonbacteraeota* than cycling and lactating females (Table S12). In particular, the genus *Helicobacter* (within the family *Epsiolonbacteroaeto*) was highly prevalent in pregnant females (Table S12), is a presumed pathogen. No predicted metabolic pathways were found to differ based on reproductive state (Table S13-S14). Age did not influence any metric of alpha diversity (Table 1 and S3, Figure S11A) or beta diversity (Table 2, Figure S11B), and no bacterial taxa (Table S6) or predicted metabolic pathways (Table S7-S8) were differentially abundant between young and old adults.

## DISCUSSION

Our findings are consistent with the hypothesis that changes in the gelada gut microbiome may help animals cope with the altered food availability and increased thermoregulatory demands associated with seasonality. First, the gelada gut microbiome was highly plastic and responded rapidly to seasonal fluctuations in climate – particularly rainfall (a proxy for available foods). Second, an increase in predicted bacterial functions involved in energy, amino acid, and lipid metabolism during both drier and colder periods suggested increased production of SCFAs, and more efficient digestion in energetically and thermoregulatory challenging periods. We further found that individual identity and social group explained nearly a third of the variation of the gelada microbiome, while other individual traits such as sex, reproductive state, and age had little effect on gut microbiome composition and function.

Rainfall was the strongest ecological factor influencing changes in the gelada gut microbiome, explaining ∼3.3% of overall microbiome composition. In particular, cellulolytic/fibrolytic and fermentative bacterial taxa increased during wetter periods when grass, which is mostly composed of cellulose, was the primary food source, while amylolytic and methanogenic bacterial taxa increased during drier periods, when geladas incorporated more starch (i.e. amylose) and lignified food into their diet. This effect of rainfall on the gut microbiome was strong, despite the fact that geladas exhibit only moderate dietary changes (i.e. from only grass to less grass and more underground organs - but from the same plant species) compared to other mammals living in more seasonal environments, e.g. that switch from ripe fruits to more folivorous diets (black howler monkeys: Amato et al. 2015; gorillas, *Gorilla gorilla gorilla* and *G. beringei* beringei: Gomez et al. 2016; Hicks et al. 2018). This pattern highlights the importance of the gut microbiome for geladas in processing their unique diet across seasons.

The efficiency of grass digestion in wet periods seems to rely on a syntropy between the first cellulolytic degraders (*Ruminococcaceae, Lachnospiraceae, Fibrobacteraceae, Spirochaetes)* and a high diversity of secondary fermenters (*Prevotellaceae* and *Bacteroidales)*, which all increase in abundance during the wet season. The first degraders attach first to the plant cell walls and hydrolyse cellulose, hemicellulose, and xylan into smaller polysaccharides and oligosaccharides (Biddle et al. 2013; White et al. 2014), while secondary fermenters ferment those soluble polysaccharides into more simple sugars (Flint et al. 2008, 2012). *Ruminococcaceae* and *Lachnospiraceae* are the two main cellulolytic taxa in mammalian gut and are commonly increasing in prevalence when animals eat more leaves and plants (Amato et al. 2015; Springer et al. 2017). In terms of secondary fermenters, *Prevotella* are widely known for their role in breaking down non-cellulosic polysaccharides and pectin (Flint et al. 2012; White et al. 2014). They are the major constituent (∼70%) of rumen bacteria (van Gylswyk 1990), and commonly increase in high fiber or fruit diets (Rampelli et al. 2015; Kovatcheva-Datchary et al. 2015; Gomez et al. 2016; Springer et al. 2017). Members of *Bacteroidales* - and particularly from the *Bacteroides* genus - have some of the largest repertoires of carbohydrate degrading activities and are able to ferment a broad range of plant polysaccharides (Salyers et al. 1977; Comstock and Coyne 2003; Flint et al. 2012; El Kaoutari et al. 2013). The increase in these cellulolytic/fibrolytic taxa and the high versatility of the secondary fermenters likely allow geladas to optimally extract nutrients from grasses eaten during wet periods.

In contrast, during drier periods, when geladas relied more on underground storage organs, we found a corresponding increase in microbial families involved in amylolytic and saccharolytic activities (*Succinivibrionaceae, Streptococcaceae, Christensenellaceae*). Interestingly, *Succinivibrionaceae* also increased during periods of energetic stress in Tibetan macaques (*Macaca thibetana*) (Sun et al. 2016) and during the dry season in the Hazda hunter gatherers of Tanzania (Smits et al. 2017), suggesting that it might help hosts cope with diet-related energy shortfalls. The gelada microbiome during the dry season was also characterized by an increase in *Methanobrevibacter*, a genus containing hydrogenotrophic archaea that converts hydrogen and formate into methane (Miller et al. 1982). The simultaneous enrichment of efficient hydrogen-producers (e.g. *Christensenellaceae (Morotomi et al. 2012), Hydrogenoanaerobacterium*: *(Song* and Dong 2009)) and formate-producers (*Succinivibrionaceae*: *(O’Herrin and Kenealy 1993)*), combined with methanogens during the dry season suggest that these taxa work together in syntropy to improve the efficiency of polysaccharide fermentation from starch in the gut in dry periods (Samuel and Gordon 2006; Basseri et al. 2010). In mice and humans, a higher abundance of methanogenic archaea was found to increase calorie harvest from diet, facilitate SCFA production by other fermentative bacteria, and stimulate lipogenesis (Samuel and Gordon 2006; Zhang et al. 2009; Basseri et al. 2010; Mathur et al. 2013).

Finally, drier periods were also characterized by a large increase in the *RFP12* family (i.e. ∼30% versus ∼18% in wetter periods) from the *Kiritimatiellaeota* phylum. The *RFP12* family remains poorly characterized but is increasingly recognized as being a keystone bacterial group in the hindgut of horses (Steelman et al. 2012; Costa et al. 2015; Edwards et al. 2020), and a common inhabitant of the rumen of sheep or cattle (Wang et al. 2017; De Mulder et al. 2017; Ribeiro et al. 2017). This suggests that it might be a keystone bacterial group for the digestion of some underground food components commonly eaten by the geladas during dry periods.

At the functional level, bacterial genes involved in energy, amino acid, and lipid metabolism increased in prevalence during the dry season. In particular, metabolic pathways linked to cellular respiration, methanogenesis, and carbon fixation pathways of prokaryotes became more common, strongly suggesting that both bacterial energy production and cellular activity were stimulated during this time. One interpretation of this data is that the increase in cellular activity simply reflects a dietary switch to starch, which is easier to hydrolyse than cellulose, and thus might more readily provoke a stimulation of bacterial activity and carbohydrate fermentation. Alternatively, the stimulation of bacterial energy metabolism and cellular activity could reflect a higher production of SCFAs by gut bacteria, supplying the host with additional energy in periods of nutrient restriction (when relying on fallback foods) (Russell and Rychlik 2001; Zhang et al. 2016). Similar increases in predicted bacterial energy metabolism have been found in energetically challenging environments (e.g. high altitude) in several other mammalian species and were correlated with higher SCFA production (Zhang et al. 2016; Li et al. 2018). Analysis of fecal SCFA profiles in geladas would help to identify if this is also the case in this high-altitude species.

While it is clear that diet shifts during drier periods, it remains unknown if (and to what extent) geladas are nutritionally or energetically constrained during this time. Grass availability declines and geladas spend more time foraging and digging for underground plant parts during the dry season (Hunter 2001; Jarvey et al. 2018). Such underground foods are usually considered fallback foods because individuals rely on them only when grass is less available and because they require long processing times (Venkataraman et al. 2014; Jarvey et al. 2018). However, one study (Hunter 2001) found that geladas obtain just as much, or even more, calories from underground storage organs as they do from grass. Whether this increased caloric intake is offset by increased foraging costs is currently unknown. However, even if increased foraging costs were demonstrated, our data suggests that the gut microbiota may increase digestive efficiency from starchy food and thereby help geladas maintain or improve energetic status during the dry season. Future studies on seasonal changes in energy balance will help resolve this issue.

In contrast to the effect of rainfall, we found mixed evidence for the effect of temperature on the gut microbiome. Temperature only explained ∼0.33% of variation in the gelada gut microbiome composition. Furthermore, few taxa shifted in abundance between the coldest and hottest months, and most taxa affected by temperature were also affected by rainfall (although the reverse was not true). This might be explained by the fact that rainfall (and thus diet) still covary with temperature to some extent (Pearson’s correlation coefficient = 0.20): geladas rely the most on underground foods in the hot-dry season (Feb to May) and the most on grass on the cold-wet season (Jun to Sep) (Jarvey et al. 2018). The cold-dry season (Oct to Jan), however, displays a mixed pattern of diet and temperature: grass availability is still high in Oct-Nov (following the rainy season) but decreases markedly in Dec-Jan (Jarvey et al. 2018; Tinsley Johnson et al. 2018). These two months are thus characterized by the introduction of underground foods in the diet and are also incidentally the coldest months of the year, making them likely the most challenging times for geladas (compounding nutritional and thermoregulatory challenges). Accordingly, cold periods were characterized by an increase in two amylolytic and lactate-producing taxa (*Streptococcus, Lactobacillus*), presumably to more efficiently extract starch from the underground foods. At the functional level, the energy and lipid metabolism of bacteria were also stimulated in the cold months, further suggesting some role of gut bacteria in stimulating host digestive efficiency and energy metabolism during thermoregulatory-demanding times.

These seasonal changes that increase energy production during colder periods may come at some cost. Such trade-offs have been proposed where shifts that benefit one aspect of host physiology consequently lead to a decrease in other microbes that may also be necessary for the host. For example, microbes that promote host digestive efficiency and energy metabolism may also promote inflammation or even suppress immune function (Vijendravarma et al. 2015; Reese and Kearney 2019). We did not detect any obvious evidence of these tradeoffs in geladas, but future work that incorporates detailed host immunological and functional microbial data is needed to help determine if such trade-offs exist.

Finally, the present study found that the gelada gut microbiome was largely explained by individual identity (20%), a pattern consistent with data from a range of vertebrates (Bik et al. 2016; Antwis et al. 2018; Trosvik et al. 2018; Kolodny et al. 2019), including humans (Costello et al. 2009; Human Microbiome Project Consortium 2012). However, the effect of social group was lower in geladas than reported for other social mammals (geladas: 6.0% vs. e.g. yellow baboon, *Papio cynocephalus*: 18.6% of variation explained (Tung et al. 2015), black howler monkey: 14% (Amato et al. 2017), ring-tailed lemurs, *Lemur catta*: 21 % (Bennett et al. 2016), Welsh Mountain ponies: 14%: (Antwis et al. 2018)). The combination of large individual effects with weak unit effects closely resembles data reported for the Guassa gelada population (Trosvik et al. 2018), suggesting a general, but consistent gelada pattern. The weak unit-level effects may result from the unique social system of geladas: because social units often aggregate into large bands whose composition change regularly, geladas may be characterized by a higher rate of inter-unit microbial transmission compared with other primates. Future studies should explore in more detail the intra-individual fluctuation in gut microbiome composition, and whether group differences in ranging patterns may explain these differences.

Other individual predictors, namely age, sex, and female reproductive state, had a very limited effect on the gut microbiome, mirroring results in other mammals (yellow baboons: (Tung et al. 2015; Ren et al. 2016), ring-tailed lemurs: (Bennett et al. 2016), Verreaux’s sifakas, *Propithecus verreauxi*: (Springer et al. 2017), chimpanzees, *Pan troglodytes schweinfurthii*: (Degnan et al. 2012), rhesus monkeys, *Macaca mulatta*: (Adriansjach et al. 2020), Welsh Mountain ponies (Antwis et al. 2018), domestic dog, *Canis lupus familiaris*: (Mizukami et al. 2019), but see black howler monkeys: (Amato et al. 2014) or Egyptian fruit bats, *Rousettus aegyptiacus*: (Kolodny et al. 2019)). Although female geladas harbored higher microbial richness than males, this resulted in minimal differences in gut microbial composition and predicted function. Compared to males, females had higher abundance of *Proteobacteria* and *Lactobacillus*. These two bacterial taxa that were previously reported to increase during pregnancy and lactation in humans and primates (Koren et al. 2012; Mallott and Amato 2018), and that act as early colonizers of the infant gut (Matsumiya et al. 2002; Martín et al. 2007; Shin et al. 2015). Additionally, pregnant female geladas harbored more *Helicobacter*, a potentially pathogenic genus (Chichlowski et al. 2008; Gao et al. 2018). An increase in potentially pathogenic microbes in pregnant females was also observed in black howler monkeys (Amato et al. 2014) and was hypothesized to be the consequence of a trade-off between reproduction and immunity. These dynamics warrant further investigation.

Overall, the gut microbiome of geladas seems to be highly plastic and can respond rapidly to changes in host diet and thermoregulatory demands. Stimulation of bacteria cellular activity could allow geladas to maintain adequate or even improved energetic balance during dry and cold periods. Our study adds to an increasing body of literature suggesting that the gut microbiota is an important system providing dietary and metabolic flexibility for the host and might be a key factor influencing the acclimatization to changing environments (Candela et al. 2012; Alberdi et al. 2016; Macke et al. 2017). In addition to fostering phenotypic plasticity, the gut microbiome is increasingly hypothesized to contribute to host evolution and speciation (Amato 2016; Alberdi et al. 2016; Macke et al. 2017) given the strong host phylogenetic signal in mammalian microbiome composition and function (Groussin et al. 2017; Amato et al. 2019) and evidence of microbiome heritability (Goodrich et al. 2014; Blekhman et al. 2015; Waters and Ley 2019). To the extent that microbiomes affect host phenotypes under selection, they will also affect host evolutionary trajectories. In the case of geladas, a shift in gut microbiome composition was probably an important adaptive mechanism that allowed members of the *Theropithecus* genus to adopt a specialized dietary niche and diversify rapidly from *Papio* ∼5 million years ago (Jablonski 2005). Contrary to host adaptive genetic mutations, which occur over the course of many generations, the gut microbiota can shift in response to changes in host diet in a matter of days (David et al. 2014). Given that the common ancestor of *Theropithecus* and *Papio* was omnivorous (Jolly 1970; Dunbar 1976), dietary flexibility provided by the gut microbiome may have been an important first step allowing members of *Theropithecus* to exploit new grassland habitats in East Africa, leading to the evolution of a specialized diet and, ultimately, further genetic and phenotypic adaptation. Future research in geladas and other mammals with peculiar dietary adaptations will further uncover how the gut microbiota influences host ecology, fitness, and the evolution of wild animal populations, and determine how an adaptable and heritable microbial community might have played a key role in supporting expansion into new habitats.

## MATERIAL & METHODS

### Study population and fecal sample collection

We collected fecal samples from a wild population of geladas living in the Simien Mountains National Park, in northern Ethiopia (13°15′N, 38°00’E). Samples were collected over a four-year period between Jan 2015 and Feb 2019. Geladas live in multi-level societies, where reproductive units (comprising a leader male, several adult females, their offspring and occasionally 1–2 follower males) and bachelor groups (comprising between 1-10 young adult males) form the smallest levels of the society, that forage and sleep together in a “band” sharing the same homerange (Snyder-Mackler et al. 2012). Since Jan 2006, the Simien Mountains Gelada Research Project (SMGRP) has collected demographic and behavioral data on over 200 individuals from two bands. All individuals are habituated to human observers on foot and are individually recognizable. Dates of birth of individuals were established using a combination of known (N=42) and estimated (N=89) birth dates. Estimated birth dates were calculated by using the mean individual age at major life-history milestones in our population (e.g. sexual maturation or first birth for females and canine eruption for males) (Beehner et al. 2009; Roberts et al. 2017). Birth dates of unknown immigrant males were estimated using an established protocol based on body size and other age-related morphological characteristics (Beehner et al. 2009). Here, we focused only on samples from adult males and females. Adult males were included when they reached 7 years of age. At this age, males have reached adult body size in stature but not in weight (Beehner et al. 2009; Lu et al. 2016), and most males have dispersed into a non-natal group (i.e. 96% of our male samples, males could thus be leaders, followers, bachelors or natals). Adult females were included after they had experienced their first sex skin swelling, a marker of reproductive maturation (which is around 4.65 years old in our population (Roberts et al. 2017)).

Fecal samples of known adult and subadult male and female subjects were collected regularly and opportunistically during the study period. Immediately upon defecation, approximately 1.5 g of feces was collected in 3 ml of RNA later (Vlčková et al. 2012; Blekhman et al. 2016), stored at room temperature for up to two months, and subsequently shipped to the University of Washington (UW). At UW, samples were stored at -80°C until the sequencing libraries were prepared. A total of 758 samples (620 female samples, 138 male samples) were collected from 131 individuals (83 females, 48 males) (mean±SD=5.79±6.14 samples per individual, range=1-21) from 28 reproductive units and 4 bachelors groups (mean±SD= 4.69±2.97 number of individuals sampled per unit, range=1-11).

The reproductive state of females at the date of sample collection was assigned based on daily monitoring of individuals for the status of sex skin swellings and the birth of infants. We assigned the three reproductive states as follows: (1) Cycling began at the first sign of postpartum sex skin swelling and ended when a female conceived - with conception defined as 183 days (mean gestation length) before the birth of a subsequent infant (Roberts et al. 2017). (2) Pregnancy started on the date of conception and ended the day before parturition. (3) Finally, lactation started on the day of parturition and ended the day before the female’s first postpartum swelling. Lactating females were further categorized as being in early lactation (infant <1 year old) or late lactation (infant >1 year old). When testing the effect of reproductive state, late lactating females were removed from the lactating category to include only females that were still nursing at the time of sample collection (females resume cycling when infants are ∼1.5 year old in our population, which is presumably accompanied by infant weaning around the same time (Roberts et al. 2017)). Furthermore, because pregnant females can abort their fetus during male takeover of their reproductive unit (Roberts et al. 2012), some pregnancies might have been misidentified as cycling based on our method of back-calculating from the date of birth. We therefore removed cycling females that experienced a takeover in the previous 6 months before the date of sample collection (N=55 samples) to avoid any misclassification of reproductive state in our analyses.

### Study site and climatic data

The study area is located at 3200m above sea level and is characterized as an Afroalpine grassland ecosystem, consisting of grassland plateaus, scrublands, and Ericaceous forests (Puff and Nemomissa 2005). Fecal samples were collected across the year, with roughly equal coverage across seasons (244 in cold-dry, 298 in cold-wet and 216 in hot-dry season as defined above). As part of the long-term monitoring of the SMGRP, daily cumulative rainfall and minimum and maximum temperature are recorded on a near-daily basis. We used the total cumulative rainfall over the 30 days prior to the date of fecal sample collection as a proxy for grass availability at the time of sample collection (Jarvey et al. 2018). In addition, we used the average minimum daily temperatures in the 30 days preceding the date of sample collection as a proxy of thermoregulatory constraints. The average minimum temperature is less correlated with cumulative monthly rainfall than the average maximum temperature in the previous 30 days (correlation coefficient: 0.25 versus -0.56) and, more importantly, is more likely to reflect the physiological effect of thermoregulation on the body (Beehner and McCann 2008; Tinsley Johnson et al. 2018).

### DNA extraction, sequencing, and data processing

We prepared 16S sequencing libraries using the protocols developed and optimized by the Earth Microbiome Project and the University of Minnesota Genomics Core (UMGC; (Gohl et al. 2016)). We extracted microbial DNA from the fecal samples using Qiagen’s PowerLyzer PowerSoil DNA Isolation kit (Qiagen #12855) following the standard protocol. We amplified the hypervariable V4 region of the 16S rRNA gene using PCR primer set 515F (TCGTCGGCAGCGTCAGATGTGTATAAGAGACAGGTGYCAGCMGCCGCGGTAA) and 806R (GTCTCGTGGGCTCGGAGATGTGTATAAGAGACAGGGACTACNVGGGTWTCTAAT) from The Human Microbiome Project and a dual**-**indexing approach (Gohl et al. 2016). Details of the amplification protocol can be accessed at https://smack-lab.com/protocols/. The first PCR round aimed at amplifying the V4 region. Each 25 μl PCR reaction well consisted of 12.5 μl of Nebnext Ultra II Q5 mastermix, 1.0 μl of each primer, and 25 ng of total DNA in 10.5 μl of nuclease-free water. PCR was performed in an Eppendorf thermocycler with a 100°C heated lid using the following cycling steps: an initial denaturing for 5 min at 95°C; followed by 15 cycles of 20 s at 98°C, 15 s at 62°C, 60 s at 72°C; and a final hold at 4°C. We cleaned up the PCR reaction with a 2:1 ratio of SPRI beads to PCR amplified DNA. The second PCR round aimed at adding a unique index primer combination to molecularly barcode each sample. We took 4 μl of product from the first PCR and added 6 μl of Nebnext Ultra II Q5 mastermix and 1 μl of of n5 and n7 indexing primers, with each sample being assigned a unique n5/n7 index primer combination. This 12 μl reaction was placed in an Eppendorf thermocycler with a 100°C heated lid, denatured for 5 min at 95°C, and amplified with 10 cycles of 20 s at 98°C, 15 s at 55°C, and 60 s at 72°C with a final hold at 4°C. After a 2:1 SPRI bead clean-up, amplification of the V4 region was confirmed in a few samples using an AATI fragment analyzer, and all libraries were quantified using a qubit fluorometer. The libraries were then pooled in roughly equimolar amounts (each with their own unique indexing primer combination), spiked with 10% PhiX to increase library complexity, and sequenced together on a single Illlumina NovaSeq 6000 SP 250 bp paired-end sequence flowcell.

We analyzed the resulting data using the Quantitative Insights Into Microbial Ecology 2 (QIIME2) platform (Caporaso et al. 2010; Hall and Beiko 2018). After trimming low quality bases from the de-multiplexed reads, we merged overlapping paired-end reads, and denoised the sequencing data by filtering and correcting Illumina amplicon sequencing errors using the Divisive Amplicon Denoising Algorithm 2 (DADA2: (Callahan et al. 2016)) plugin incorporated in QIIME2. DADA2 infers sequences exactly resulting in amplicon sequence variants (ASVs). Forward and reverse reads were trimmed to 220 and 180 bases, respectively, to remove the low**-** quality portion of the sequences. The forward and reverse reads were then merged together and chimeric sequences were removed. Only samples with more than 20,000 reads were retained for analysis (following observation of rarefaction curves, Figure S12). After filtering, trimming, merging, and chimera removal, we retained a total of 348,390,395 reads across the 758 fecal samples (459,618±815,020 reads per sample, range=20,109-10,735,588). ASVs were taxonomically assigned using the q2**-**feature classifier in QIIME2 against version 132 of the SILVA database (updated December 2017) (Quast et al. 2013) based on 100% similarity. Uninformative taxonomic assignments of ASVs found in SILVA (e.g. “wallaby metagenome”, “unassigned bacteria”, etc.) were converted to “NA” to simplify analysis at higher taxonomic levels. All ASVs belonging to the order *WCHB1-41* (phylum *Kiritimatiellaeota*) were not assigned at the family level in the SILVA classification. However, in the Greengene classification (version 13_8) all ASVs from this order in the gelada gut were assigned to the *RFP12* family. Thus, we attribute the family *RFP12* to all ASVs from the order *WCHB1-41* in SILVA classification.

### Statistical analyses

The count and taxonomy files generated by QIIME2 were imported into R version 3.5.2 (Team and Others 2013) using the qiime2R package (Bisanz 2008) and analyzed using the phyloseq package (McMurdie and Holmes 2013). The majority of the 19,606 ASVs in our dataset were found at very low frequency or only in one sample (71% of ASVs were found in only one sample and 6.2% of ASVs were not assigned at the phylum level). Thus, we further filtered the count table to retain only ASVs that had at least 500 reads in total in the dataset (i.e. 0.00014% relative abundance) to eliminate potentially artifactual sequences. With this filtering criteria, only 3,295 ASVs remained, with all of them assigned at the phylum level and most (97%) observed in at least two samples (Figure S13). Most ASVs could be taxonomically assigned to the class and order levels (∼99%), with assignments decreasing at the family (85%) and genus (61%) levels.

The use of rarefaction (i.e. subsampling of the read count in each sample to a common sequencing depth) has been discouraged due to the loss of information and precision (McMurdie and Holmes 2014), as well as the use of count normalization methods from the RNA-seq field (e.g. DESeq2 or edgeR). However, microbiome datasets are more sparse (zero-inflated) and more asymmetrical than genetic expression datasets (Gloor et al. 2017; Weiss et al. 2017). Thus, we used a compositional approach when possible (e.g. centered-log-ratio normalization of the counts and Aitchison distance for beta diversity analysis) (Gloor et al. 2016, 2017), controlling for sample sequencing depth in multivariate analyses to account for repeated samples from the same individual.

We replicated alpha- and beta-diversity analyses using traditional rarefaction methods to facilitate comparisons with other studies. To generate the rarefied dataset, we randomly sampled 20,000 reads from the raw fastq files of each sample and processed this new rarefied dataset into the DADA2 pipeline. This dataset was further filtered to remove the low frequency ASVs (i.e. ASVs not included in the pool of 3,295 ASVs retained in the full dataset). This resulted in a dataset containing the same 758 samples, with 2853 ASVs and with relatively homogenous sequencing depth (18205±1415 reads per sample, range=7460-19444).

All mixed models described below were run using either the lmer (for linear mixed models, LMMs) or glmer (for binomial and negative binomial generalized linear mixed models, GLMMs) functions of the lme4 package (Bates et al., 2014). All quantitative variables (i.e. cumulative rainfall, averaged temperature, and age) were z-transformed to have a mean of zero and a standard deviation of one to facilitate model convergence. The significance of the fixed factors was tested using a likelihood ratio test, LRT (assuming an asymptotic chi-square distribution of the test statistic) via the drop1 function. To test for significant pairwise differences between levels of multilevel categorical variables (i.e. reproductive state), *post hoc* Tukey’s Honest Significant Difference tests were carried out using the multcomp package in R (Hothorn et al. 2008).

#### Alpha-diversity analyses

We calculated three measures of alpha diversity: observed richness (the total number of different ASVs in a sample), Shannon diversity index (accounts for both richness and evenness of ASVs in a sample), and Faith’s phylogenetic diversity (accounts for phylogenetic distance between bacterial species, using the picante package (Kembel et al. 2010)). We modeled each alpha diversity metric using linear mixed models: (i) as a function of age, sex, cumulative monthly rainfall, average monthly minimum temperature, and sequencing depth of the sample (N=758 samples), and (ii) as a function of reproductive state (cycling, early lactating and pregnant), age, cumulative monthly rainfall, and average monthly minimum temperature in samples collected from females (N=439). Individual identity and unit membership were included as random effects to control for individual and unit repetition across samples. We also ran the same models on the rarefied dataset (Table S15).

#### Beta-diversity analyses

We then assessed how the same predictors were associated with between-sample community dissimilarity. To account for differences in sequencing depth between samples, the counts were normalized using the centered-log-ratio (CLR) method (and using a pseudocount of 0.65 for zero counts) from the “compositions” package (van den Boogaart and Tolosana-Delgado 2008). We then calculated the Aitchison distance between samples (i.e. simply the Euclidean distance between samples after clr transformation of the counts) (Aitchison et al. 2000) and conducted a Principal Component Analysis (PCA) (function “prcomp”) to visually represent between-samples dissimilarity according to the predictors. This approach has been recommended for microbiome datasets (Gloor et al. 2017), and allows for the projection of each sample onto individual principal components (PCS) and the variable loadings of ASVs onto each PC. While the first axis of variation correlated mostly with rainfall (Figure 1C), the second PCA axis was correlated with sequencing depth, and explained 11% of the variation (Figure S14). We used Permutational Multivariate Analysis of Variance (PERMANOVA) tests to assess the effect of the predictors on the Aitchison distance between samples (using using 10,000 permutations and the “adonis2” function from the “vegan” package (Oksanen et al. 2010)). We ran three different models: (1) including all samples where we tested only the effect of individual identity and sequencing depth, (2) including all samples where we tested the effect of unit, age, sex, cumulative monthly rainfall, average monthly minimum temperature, and sequencing depth of the sample and (3) including only female samples where we tested the effect of unit, reproductive state, age, cumulative monthly rainfall and average monthly minimum temperature. In models 2 and 3, individual identity was included as a blocking factor (“strata”) to control for repeated sampling. We also replicated beta diversity analysis on the rarefied dataset. We ran PERMANOVA tests using three complementary pairwise dissimilarity metrics (Bray-Curtis distance, unweighted and weighted UniFrac distances) to assess between-sample variation according to the same predictors (the same three models). Beta diversity results remained qualitatively similar (Table S16).

#### Differential abundance testing

We examined how our predictors were associated with differential abundance of bacteria (at the phylum, class, order, family and genus levels) using negative binomial GLMMs. Compared to LMMs, negative binomial mixed models are better equipped to handle over-dispersed and zero-inflated distributions that often characterize microbiome datasets (Zhang et al. 2018). They also facilitate tests of several independent predictors while taking into account longitudinal designs including random effects. We first aggregated the counts (i.e. the number of reads per taxa and per sample) at the taxonomic level of interest. Only taxa that had an average relative abundance across samples ≥0.01% were tested. Then, for a given taxa, the count per sample was modeled as a function of: (1) age, sex, cumulative monthly rainfall and averaged monthly minimum temperature (all samples), or (2) female reproductive state, age, cumulative monthly rainfall and averaged monthly minimum temperature (female samples only). The log-transformed number of reads per sample was included as an offset term to control for variation in sequencing depth across samples. Individual identity and unit membership were included as random effects in all models. When negative binomial models failed to converge in some taxa, we converted the counts in presence/absence and modeled them with binomial GLMMs. Benjamini-Hochenberg corrected p- values < 0.05 were considered statistically significant.

#### Functional profiling of microbiota

We estimated the bacterial and archaeal genes present in the metagenomes of each sample using Phylogenetic Investigation of Communities by Reconstruction of Unobserved States version 2 (PICRUSt2) (Douglas et al. 2019). In brief, ASVs were aligned to reference sequences using HMMER (Finn et al. 2011) and placed into a reference tree using EPA-NG (Barbera et al. 2019) and gappa (Czech 2019). PICRUSt2 normalizes for multiple 16S gene copies in bacteria using castor, a hidden state prediction tool (Louca and Doebeli 2018). The normalized data were used to predict gene family profiles, and mapped onto gene pathways using MinPath (Ye and Doak 2009). We followed the default protocols outlined on the PICRUSt2 GitHub page (https://github.com/picrust/picrust2/wiki). We investigated the predicted gene families using the Kyoto Encyclopedia of Genes and Genomes (KEGG) Orthology (KO) database. The accuracy of the PICRUSt2 predictions for each sample were assessed by calculating the weighted Nearest Sequence Taxon Index (NSTI) score, a measure of how similar the bacteria from the sample are to reference genome sequences. The average NSTI value across all samples was high in geladas (mean±SD=0.60±0.13) compared to other mammals (Douglas et al. 2019), so the results of this analysis should be interpreted with caution. Five ASVs (out of 3295) had a NSTI score>2 and were removed from our final predictions. The association between the relative abundance of functional categories as estimated by PICRUSt2 and the predictors (on all samples or female samples only) were examined using LMMs. Only functional pathways that had ≥ 0.1% relative abundance across samples were tested. Individual identity and unit membership were included as random effects in all models.

## DATA AVAILABILITY

All 16S sequence data used in this study are available at the NCBI Sequence Read Archive (https://www.ncbi.nlm.nih.gov/) under BioProject ID PRJNA639843. Data and code (including how to run the QIIME2 pipeline on our data) are available at: https://doi.org/10.5281/zenodo.3932310.

## ACKNOWLEDGEMENTS

We thank the Ethiopian Wildlife Conservation Authority (EWCA), along with the wardens and staff of the Simien Mountains National Park for permission to conduct research and ongoing support to our long-term research project. We are also very grateful to the Simien Mountains Gelada Research Project field team for their help with field data collection, particularly our primary data collectors: Eshete Jejaw, Ambaye Fanta, Setey Girmay, Atirsaw Adugna, and Dereje Bewket. Special thanks to Johannes R. Björk and Elizabeth Archie for stimulating discussion about microbiome analyses and sharing code. This research was funded by the National Science Foundation (BCS-1723228, BCS-1723237, BCS-2010309, BCS-0715179, IOS-1255974, IOS-1854359). KRA is supported as a fellow in the CIFAR ‘Humans and the Microbiome’ program. NSM was supported by the National Institutes of Health (R00-AG051764). The long-term gelada research was supported by the University of Michigan, Stony Brook University, and Arizona State University.

## AUTHOR CONTRIBUTIONS

Conceptualization, Data curation, Formal analysis, Investigation, Visualization: A.B, A.L., N.S.M; Methodology: A.B., S.S., A.M., R.P.; Writing – Original Draft: A.B, A.L., N.S.M; Writing – Review & Editing: all authors; Funding Acquisition: A.L.,N.S.M.,J.C.B., T.J.B.,L.R.; Supervision: A.L. and N.S.M.

## COMPETING INTERESTS

The authors declare that they have no competing interests.

## Notes

https://doi.org/10.5281/zenodo.3932310

### Competing Interest Statement

The authors have declared no competing interest.

