## Supplemental Figures for "Seasonal shifts in the gut microbiome indicate plastic responses to diet in wild geladas"

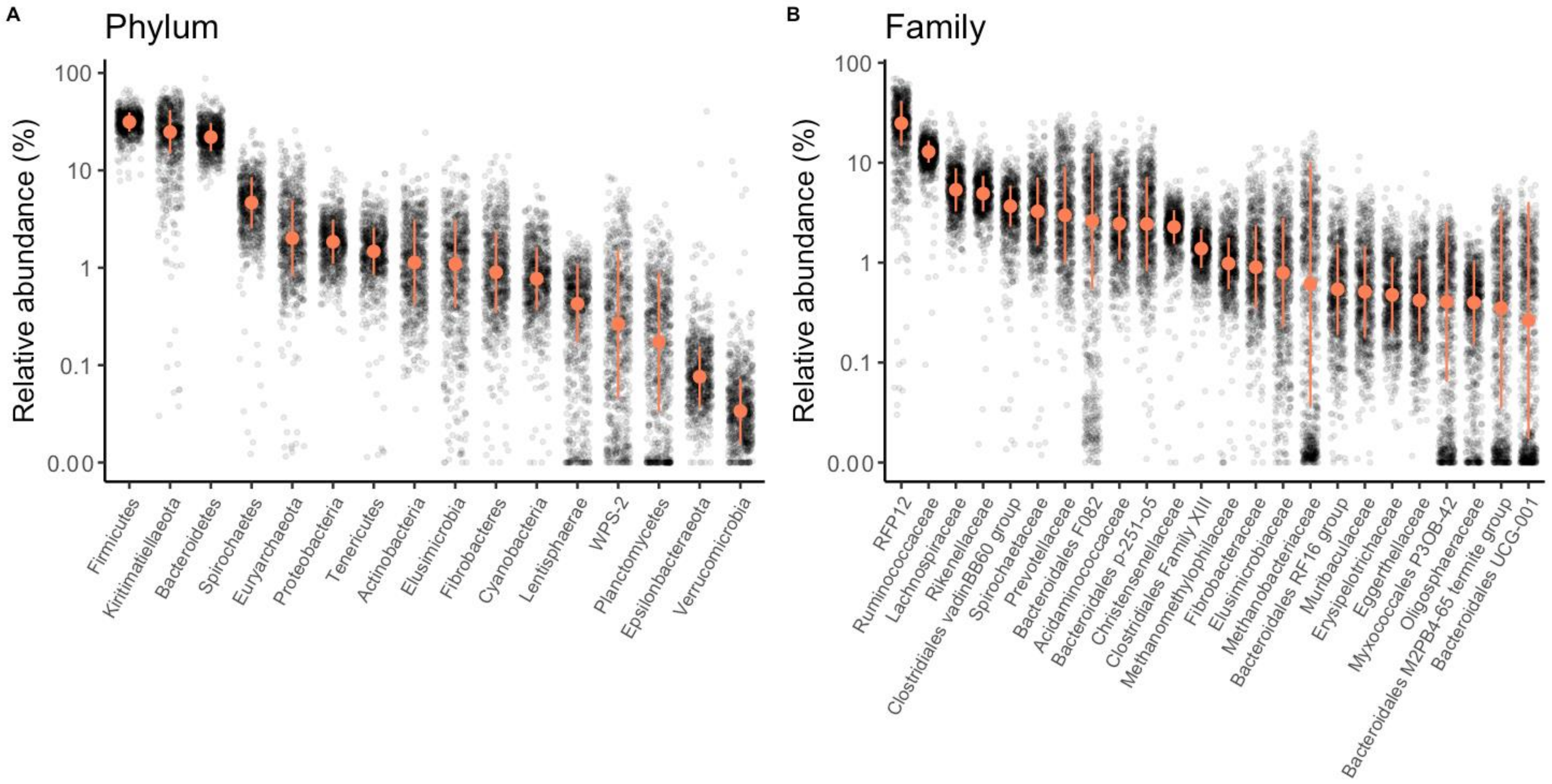


**Supplemental Figure 1. Taxonomic composition of the geladas gut at the phylum and family levels.** Relative abundance **(A)** of all bacterial phyla and **(B)** of the 24 most abundant families (whose relative abundance>0.02%) in the gelada feces. Compared to Figure 1, the tick marks on the y-axis are spaced on a log10 scale to better represent the variation across samples. The median and median absolute deviation (error limit) are represented in orange.

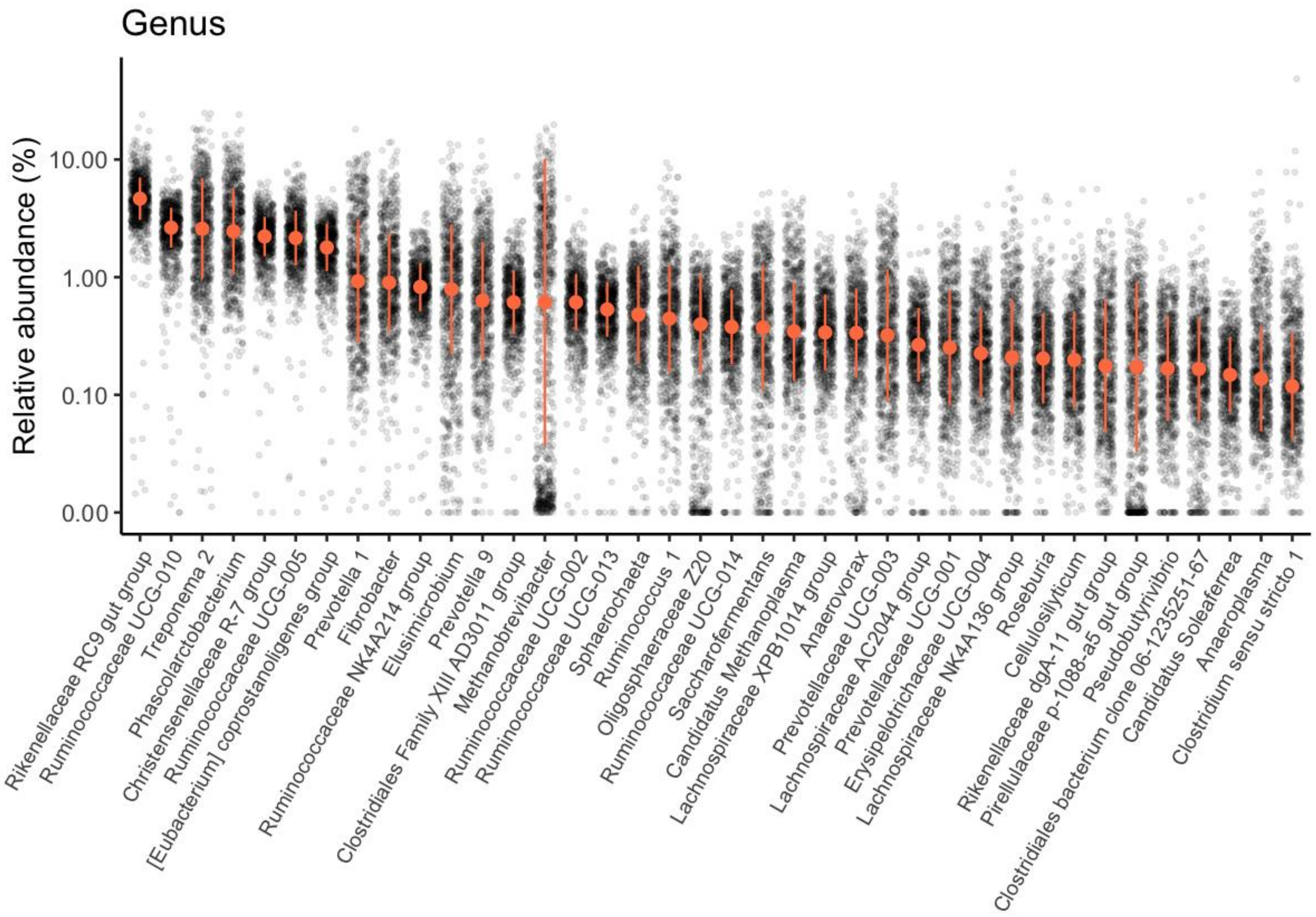


**Supplemental Figure 2. Genus composition of the geladas gut.** Relative abundance of the 38 most abundant genera (whose relative abundance>0.01%) in the gelada feces. The tick marks on the y-axis are spaced on a log10 scale. The median and median absolute deviation (error limit) are represented in orange.

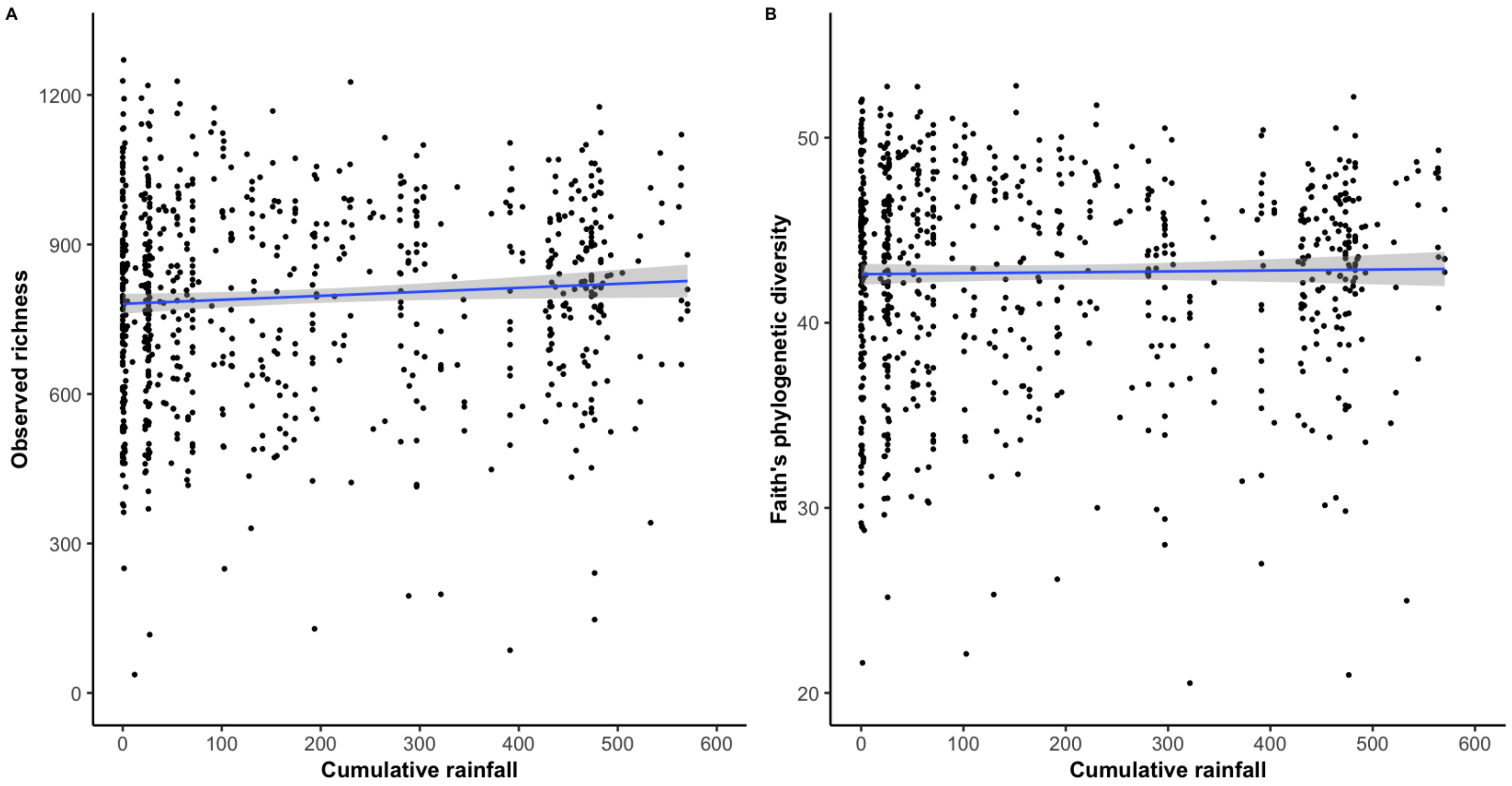


**Supplemental Figure 3. Rainfall is not associated with Observed richness and Faith's phylogenetic diversity.** Partial residual plot of **(A)** Observed richness and **(B)** Faith's phylogenetic diversity (PD) according to cumulative rainfall (in mm). Black dots represent the partial residuals from the LMM (i.e. showing the association between cumulative rainfall and alpha diversity, while controlling for all other predictors). The blue line and confidence intervals come from a linear regression (for representation only). One and 5 outlier samples (with a particularly low diversity) were omitted for panel A and B respectively for clarity of representation.

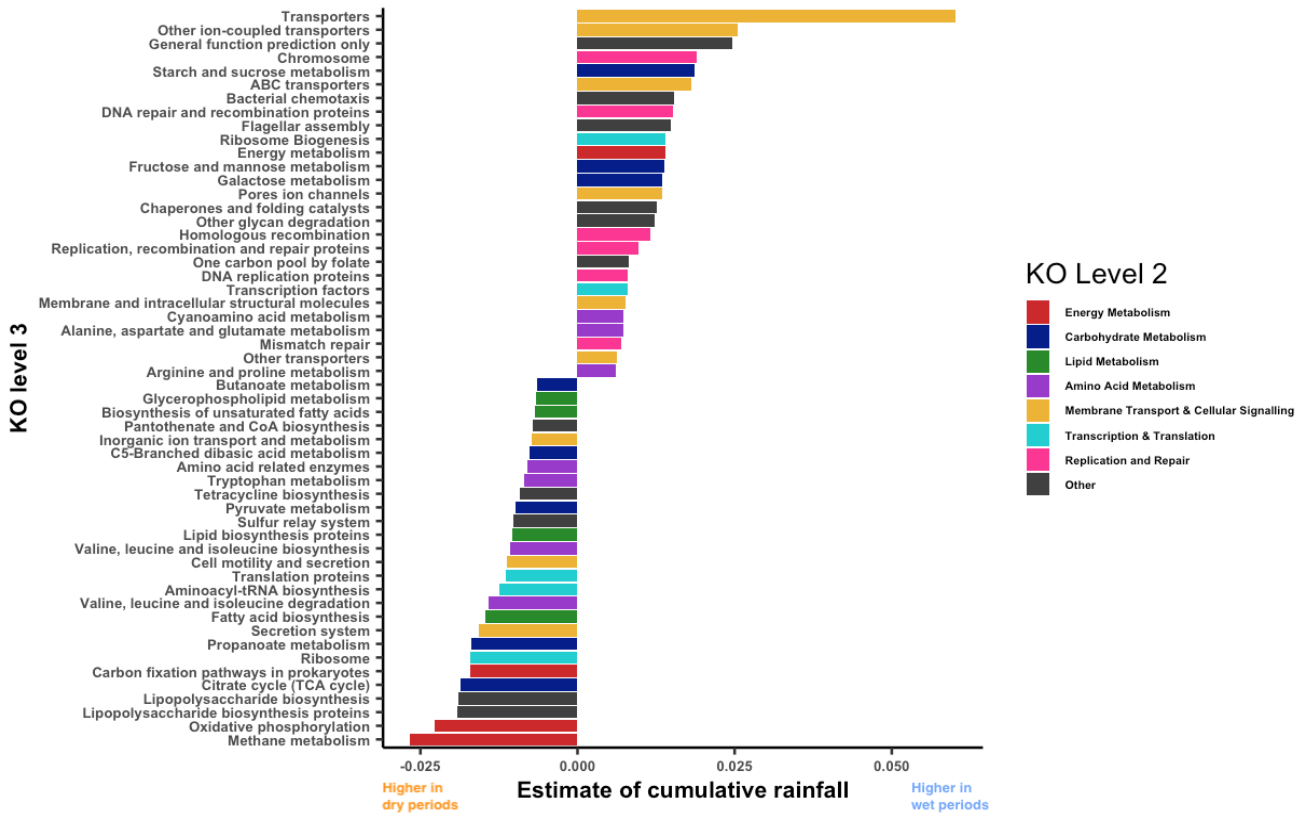


**Supplemental Figure 4. Bacterial functional pathways that significantly associated with cumulative rainfall at KO level 3.** The estimate of the cumulative rainfall effect comes from a LMM fitted on the relative abundance of each pathway per sample. Only pathways with $p_{BH} < 0.05$ were considered significant. For ease of representation, only pathways with effect sizes > |0.006| are represented. The full list can be found in Table S8. Classification of KO level 3 pathways in broader categories were based on their KO level 2 assignment, with a few changes made for clarity of representation. Level 3 pathways from *Metabolism of Other Amino Acids* (level 2) were reclassified in the "Amino Acid Metabolism", *Translation proteins* and *Replication, recombination and repair proteins* (both level 3 and initially in Genetic Information Processing at level 2) were reclassified in "Transcription & Translation" and "Replication and Repair" respectively. The category "Membrane Transport & Cellular Signalling" regroups pathways from "Membrane Transport" and the other pathways from "Cellular Processes and Signaling"

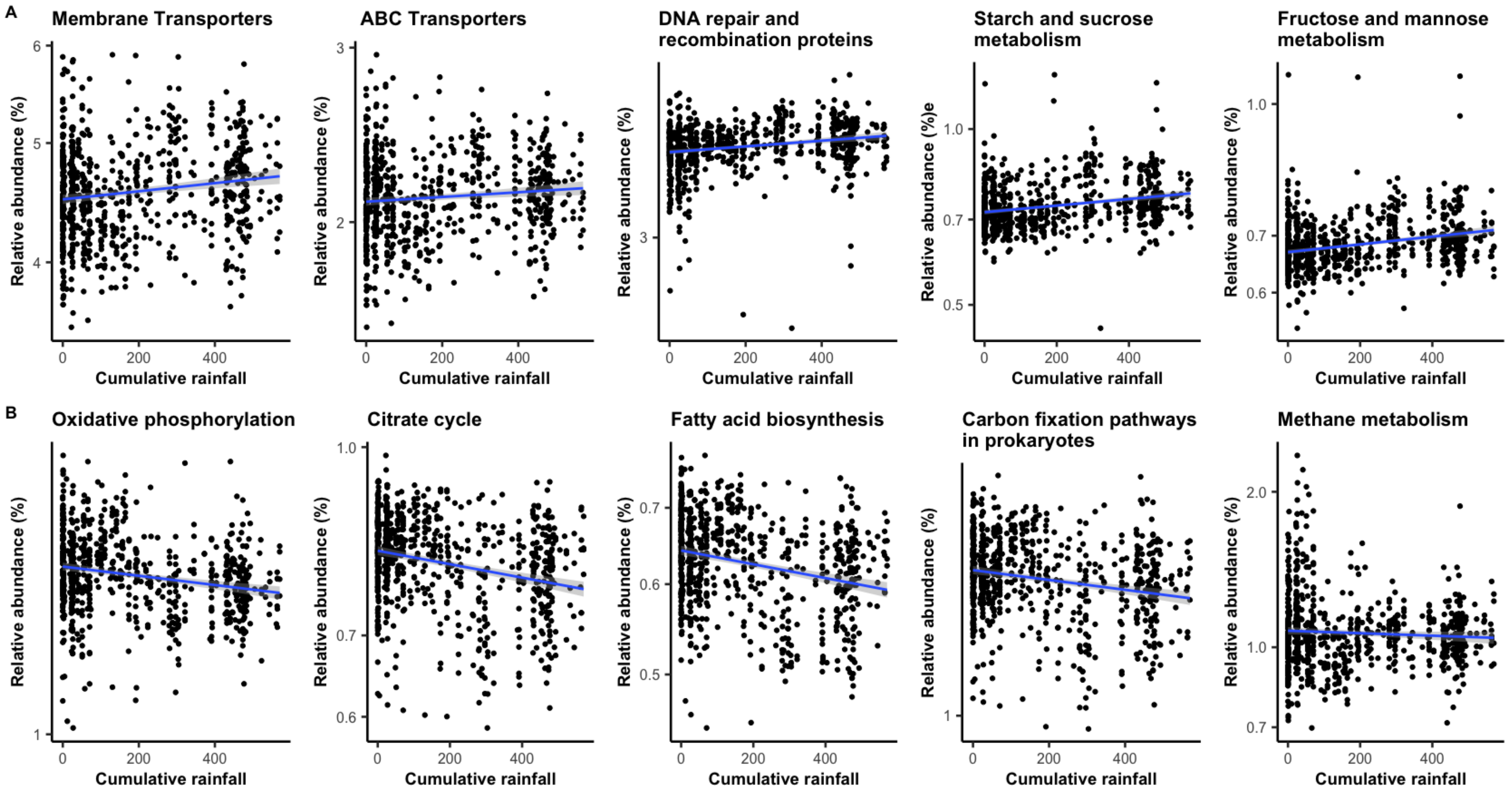


**Supplemental Figure 5. Rainfall predicts the functional profile of the gut microbiome.** Relative abundance of ten functional pathways (at KO level 3) that are enhanced **(A)** during the wet season and **(B)** during the dry season. Note that the tick marks on the y-axis are spaced on a log10 scale. The blue line and confidence intervals come from a linear regression (for representation only). The significance of the rainfall effect effects per pathway have been estimated using LMMs including individual and unit membership as random effects.

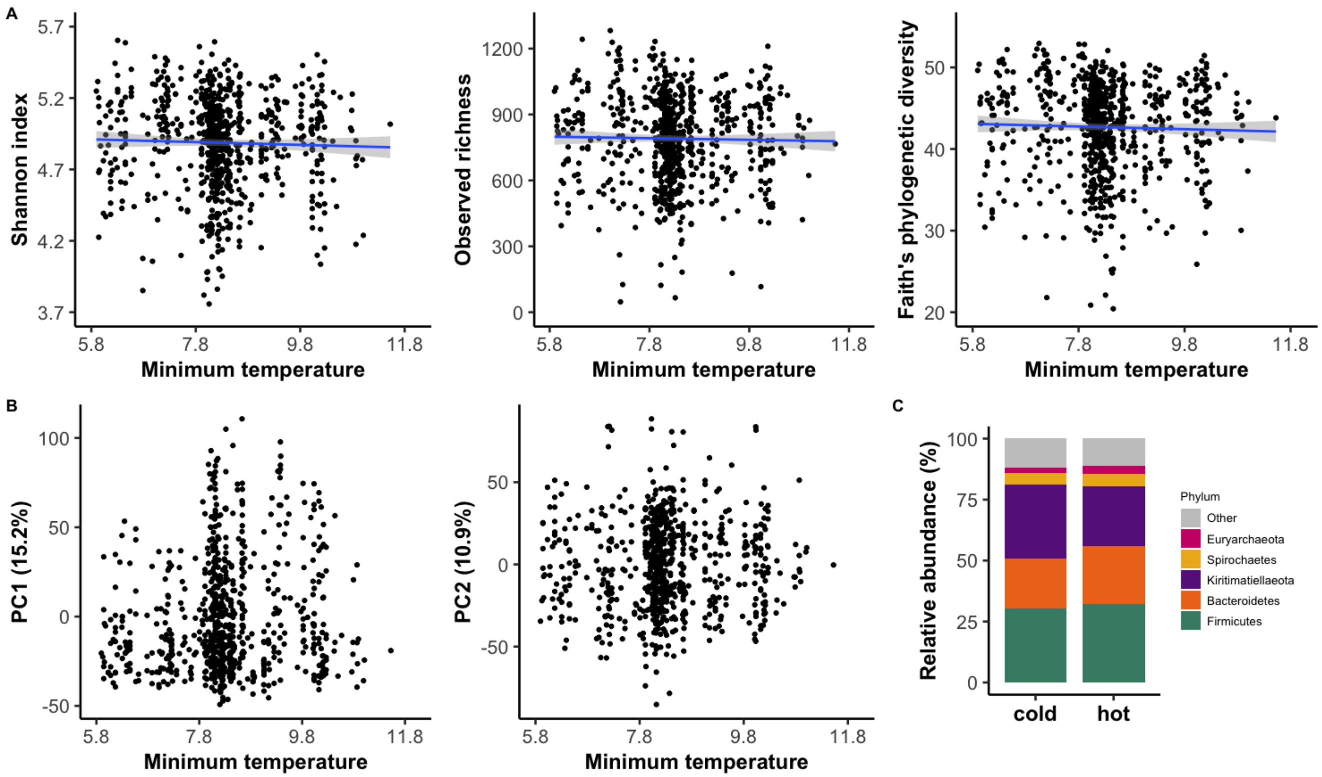


**Supplemental Figure 6. Small effect of ambient temperature on the gelada gut microbiome. (A)** Partial residual plots of the three alpha diversity indices (Shannon index, Observed richness and Faith's phylogenetic diversity) according to the average minimum temperature in the previous month of sample collection (in $^{o}$C). Black dots represent the partial residuals from the LMM (i.e. showing the association between temperature and alpha diversity, while controlling for all other predictors). The blue line and confidence intervals come from a linear regression (for representation only). Nine, 1 and 5 outlier samples (with a particularly low diversity) were omitted for Shannon, richness and Faith's PD respectively for clarity of representation. **(B)** Visualization of between-sample dissimilarity (based on Aitchison distance) on the first and second principal component according to minimum temperature. **(C)** Compositional barplot of the five most abundant phyla in the cold (i.e. <8$^{o}$C in the past month, N=191) and hot (>8$^{o}$C in the past month, N=567) samples (minimum temperature was converted to a categorical variable for representation purposes).

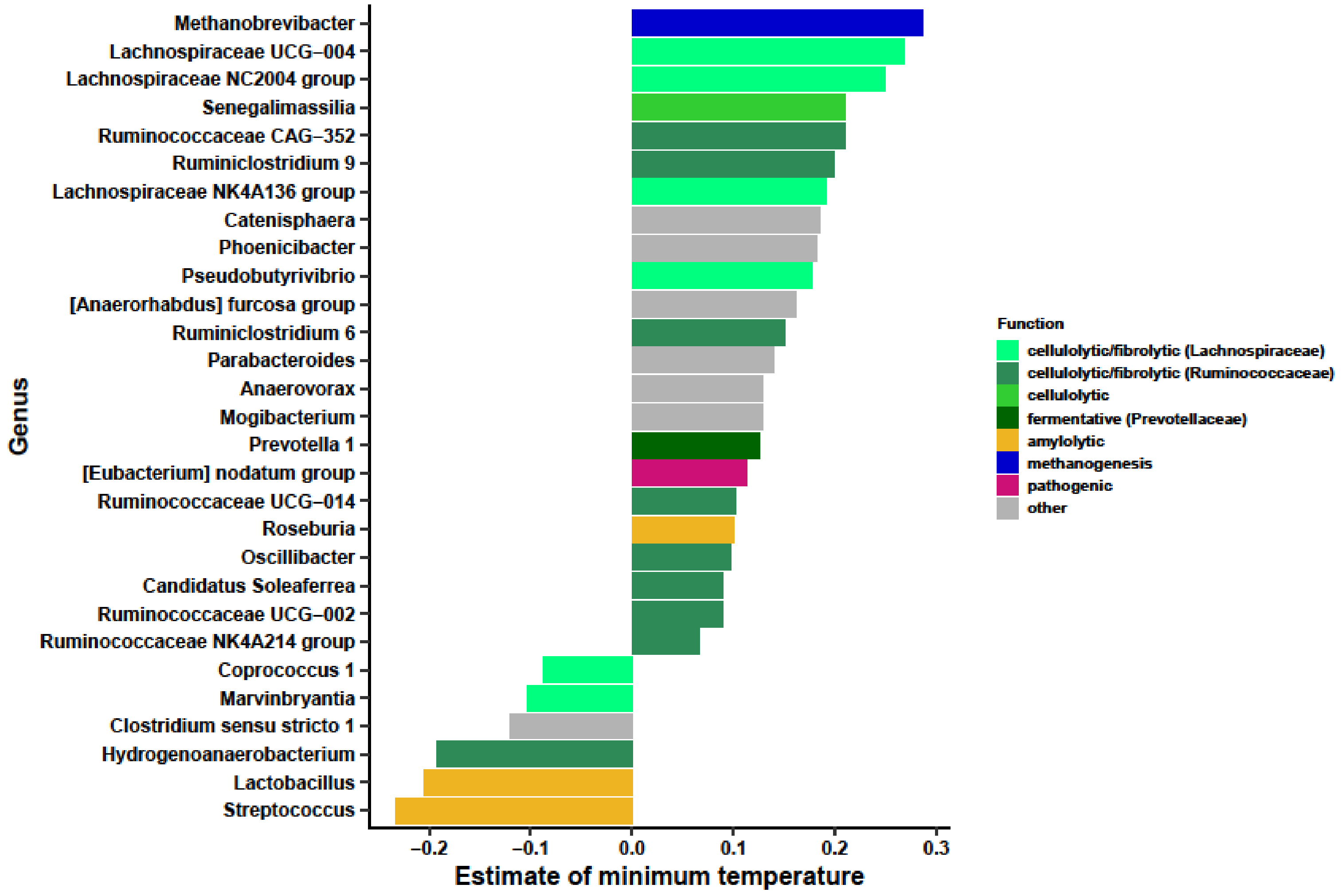


**Supplemental Figure 7. Genera that significantly associated with average minimum temperature.** The estimate of the "temperature" effect for each taxa comes from a negative binomial GLMM fitted on the count of each taxa per sample (controlling for sample sequencing depth as an offset factor, and including individual and unit membership as random effects). Only taxa with $p_{BH} < 0.05$ were considered significant. The full list can be found in Table S6.

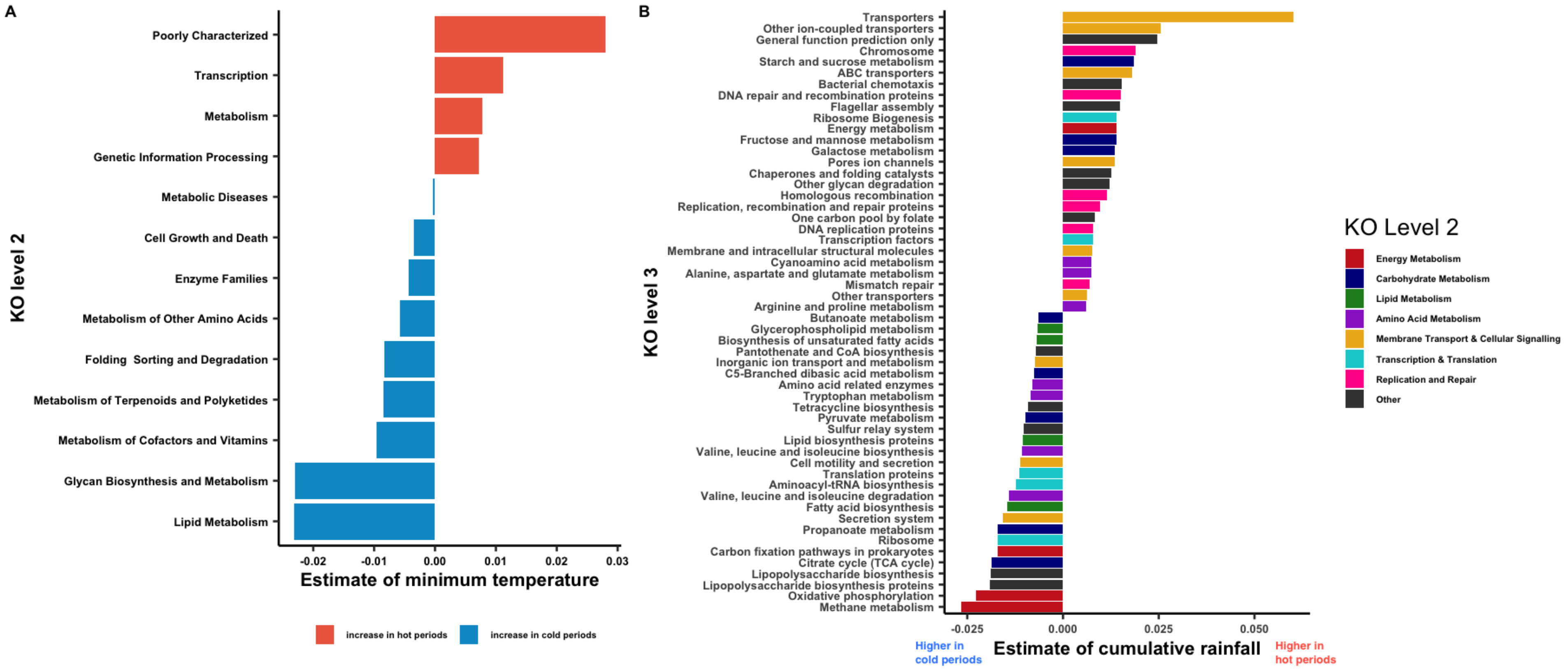


**Supplemental Figure 8. Bacterial pathways that are found differentially abundant according to averaged minimum temperature at KO (A) level 2 and (B) level 3.** The estimate comes from a LMM fitted on the relative abundance of each pathway per sample. Only pathways with $p_{BH} < 0.05$ were considered significant. For ease of representation on panel B, only pathways with effect size > |0.002| were represented. The full list can be found in Table S8.

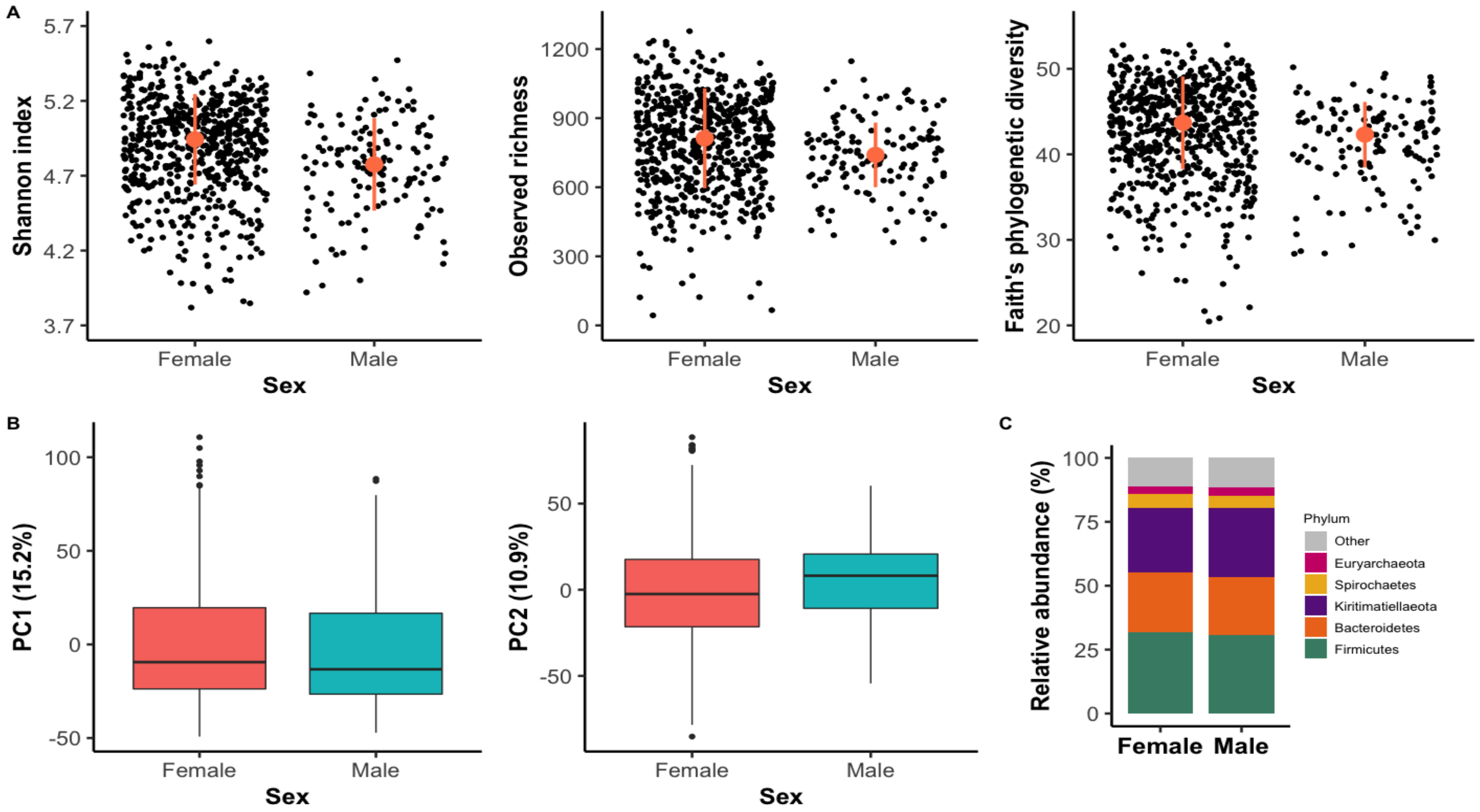


**Supplemental Figure 9. Effect of sex on the gelada gut microbiome. (A)** Partial residual plots of the three alpha diversity indices (Shannon index, Observed richness and Faith's phylogenetic diversity) according to the sex of the sampled individual. Black dots represent partial residuals of the LMM. The median and median absolute deviation (error limit) of the distribution are represented in orange. Ten, 1 and 5 outlier samples (with a particularly low diversity) were omitted for Shannon, richness and Faith's PD respectively for clarity of representation. **(B)** Visualization of between-sample dissimilarity (based on Aitchison distance) on the first and second principal component according to sex. **(C)** Compositional barplot of the five most abundant phyla in male (N=138) and female (N=620) samples.

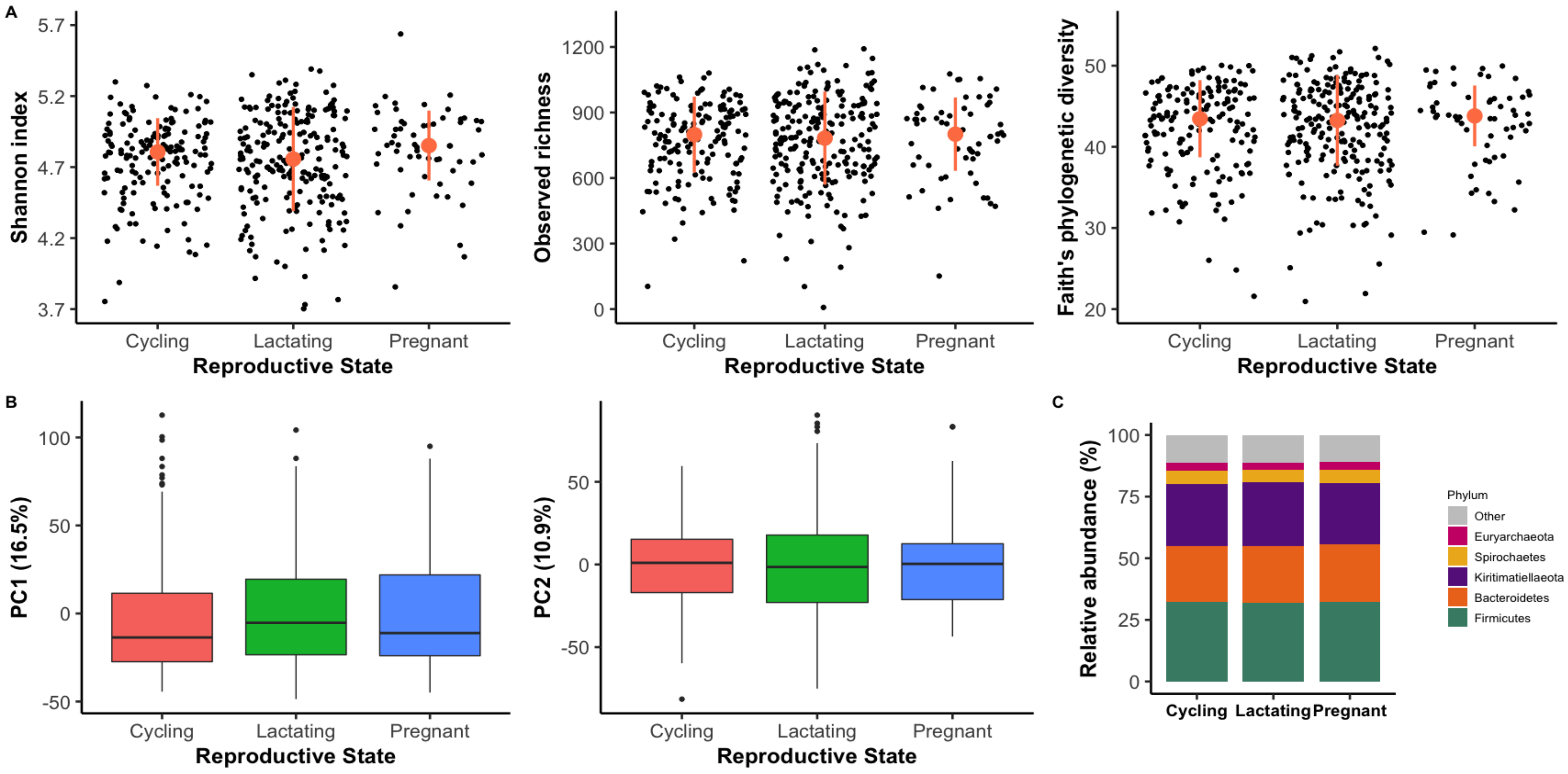


**Supplemental Figure 10. Effect of female reproductive state on the gelada gut microbiome. (A)** Partial residual plots of the three alpha diversity indices (Shannon index, Observed richness and Faith's phylogenetic diversity) according to the reproductive state of the sampled female. Black dots represent partial residuals of the LMM. The median and median absolute deviation (error limit) of the distribution are represented in orange. Ten, 1 and 5 outlier samples (with a particularly low diversity) were omitted respectively for clarity of representation. **(B)** Visualization of between-sample dissimilarity (based on Aitchison distance) on the first and second principal component according to reproductive state. **(C)** Compositional barplot of the five most abundant phyla in pregnant (N=61), lactating (N=346) and cycling (N=158) female samples.

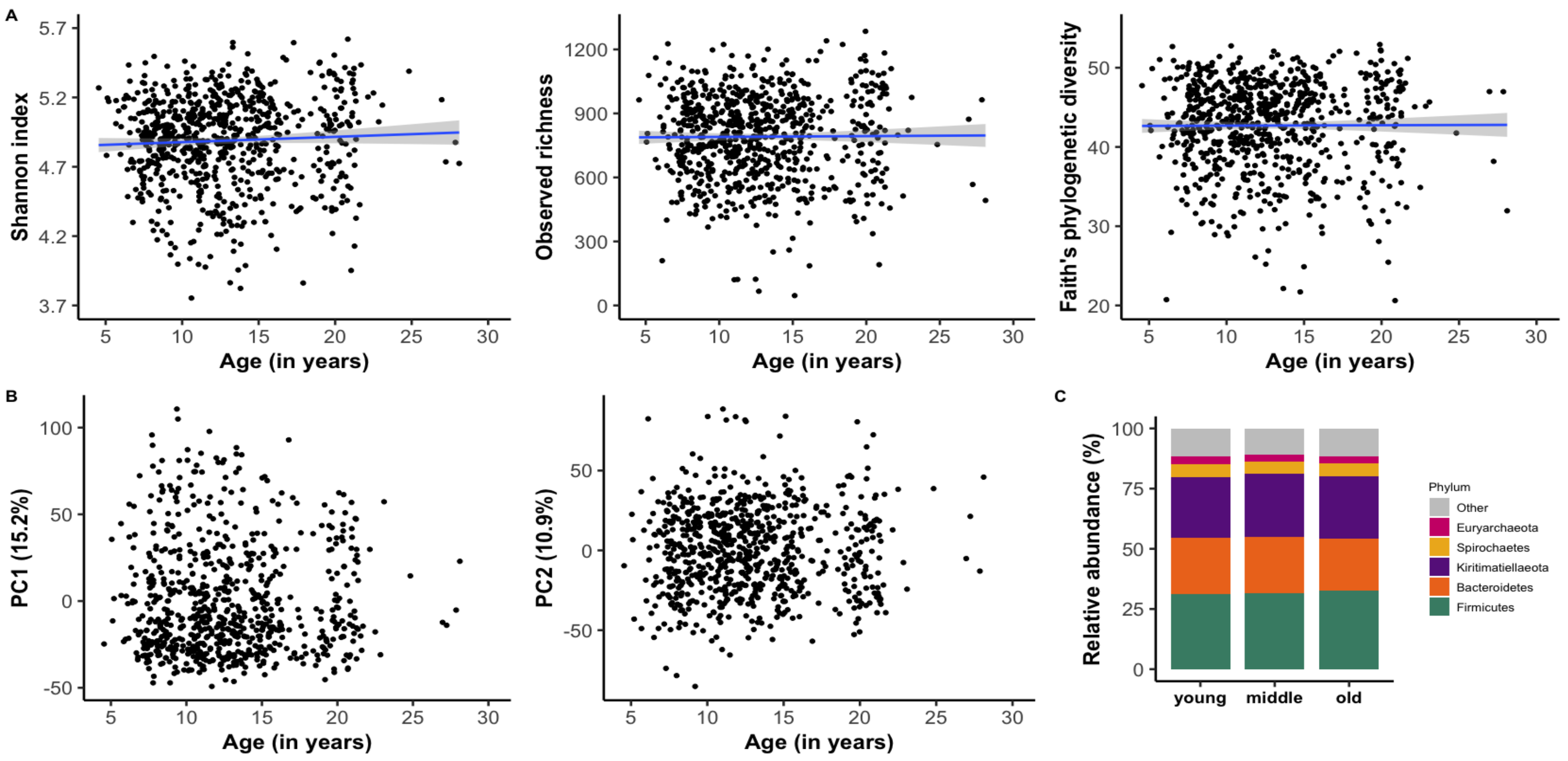


**Supplemental Figure 11. Effect of individual age on the gelada gut microbiome. (A)** Partial residual plots of the three alpha diversity indices (Shannon index, Observed richness and Faith's phylogenetic diversity) according to the age of individuals at the date of sample collection in years). Black dots represent the partial residuals from the GLMM (i.e. showing the association between age and alpha diversity, while controlling for all other predictors). The blue line and confidence intervals come from a linear regression (for representation only). Nine, 1 and 5 outlier samples (with a particularly low diversity) were omitted respectively for clarity of representation. **(B)** Visualization of between-sample dissimilarity (based on Aitchison distance) on the first and second principal component according to age. **(C)** Compositional barplot of the five most abundant phyla between young (<10 years old, N=215 samples), middle-aged (10 to 17 years old, N=420 samples) and old (>17 years old, N=123 samples) individuals (age was converted to a categorical variable for representation purposes only).

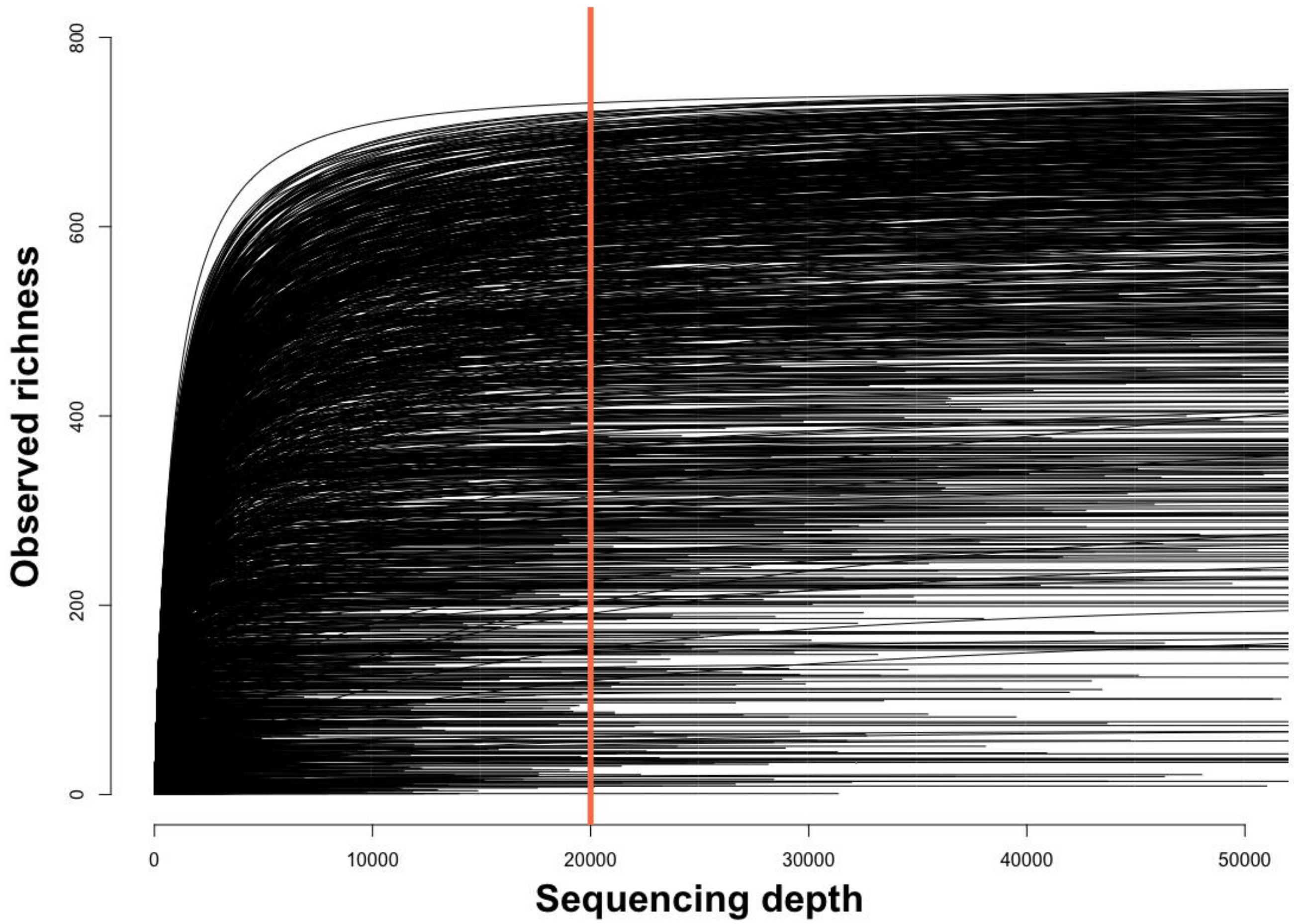


**Supplemental Figure 12.** Rarefaction curves of samples. Only samples that had at least 20000 reads were included in this study.

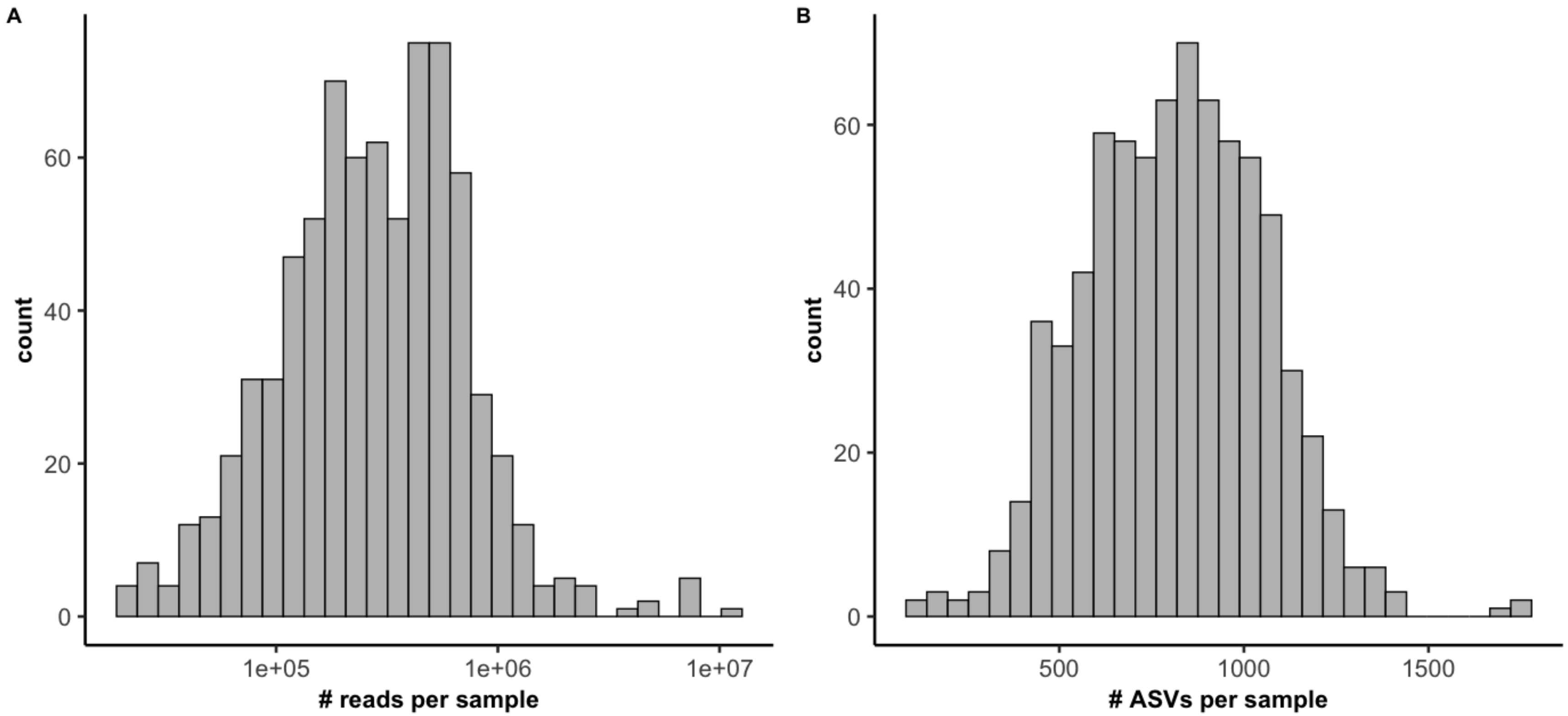


**Supplemental Figure 13. 16S sequencing and dataset characteristics. (A)** Distribution of the total number of reads per sample (the tick marks on the x-axis are spaced on a log10 scale). **(B)** Distribution of the total number of ASVs per sample.

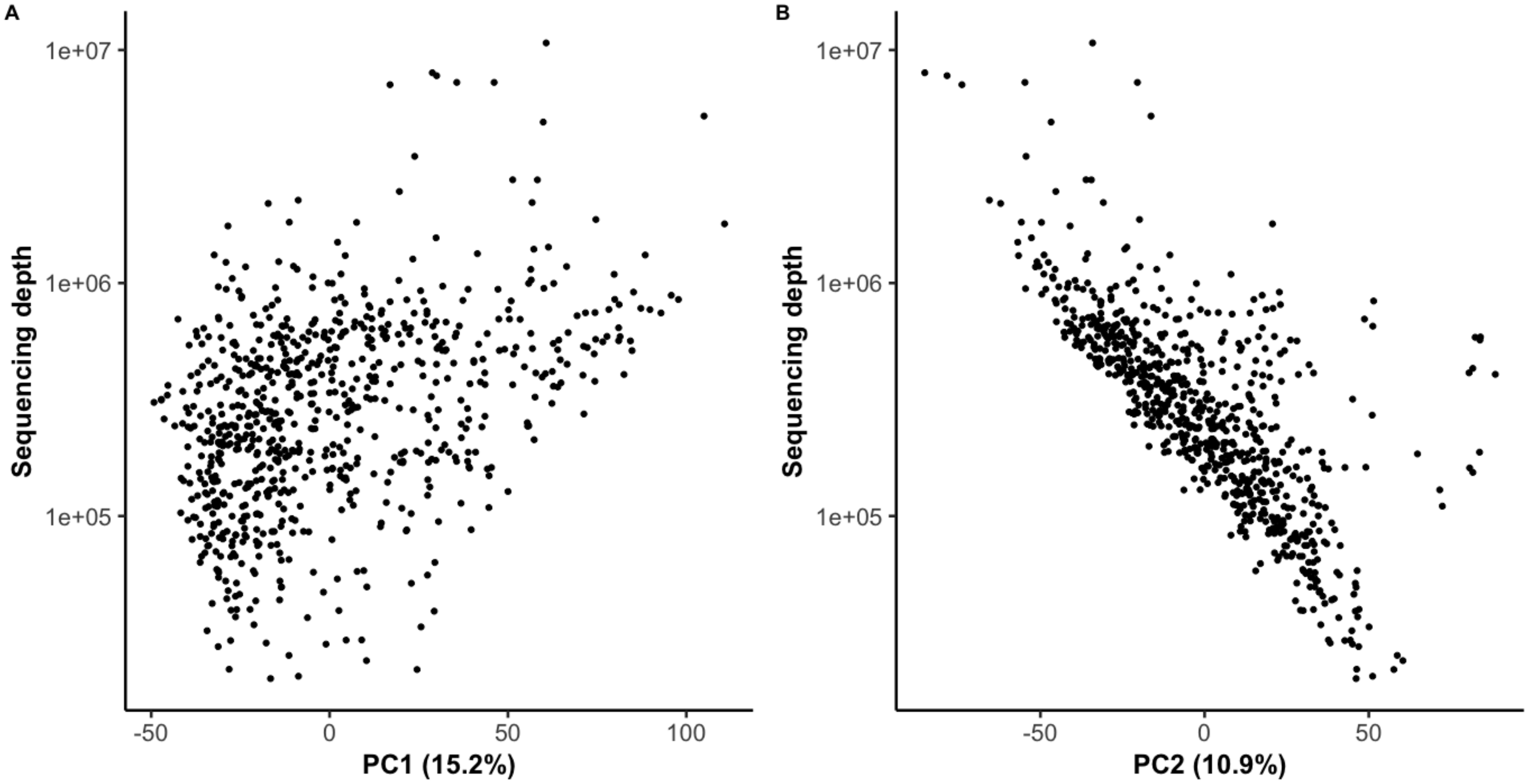


**Supplemental Figure 14.** Visualization of differences in the gut microbiome composition according to sequencing depth of the samples based on Aitchison distance dissimilarity matrix. Points represent individual samples.
