## Supplementary material for "Seasonal shifts in the gut microbiome indicate plastic responses to diet in wild geladas": Legend Supplemental Tables

**Legend of Supplementary Tables**

**Table S1. Taxonomic composition of the gelada gut at the phylum, class, order, family and genus levels.** The mean relative abundance and prevalence (% of the samples with each taxon) of bacterial taxa are indicated.

**Table S2. Taxonomic composition of the core ASVs (i.e. present in at least 90% of samples) in the gelada gut at the order level.**

**Table S3. Predictors of observed richness and Faith's phylogenetic diversity (PD).** Estimates with p-values <0.05 are highlighted in bold.

**Table S4. Loading scores of ASVs on the first principal component.** Positive loadings correspond to wetter periods, while negative loadings correspond to drier periods.

**Table S5 Taxonomic distribution of seasonally differentially abundant taxa.** Only taxa with a PC loading scores > 0.4 and <-0.4 are included.

**Table S6. Differential abundance results for all taxa.** Results were obtained by fitting negative binomial GLMMs for each taxa, controlling for individual identity and unit membership. The estimate and p-values of fixed effects are reported. Due to some negative binomial models that did not converge, taxa preceded by a "*" were modeled with a binomial GLMM, with presence/absence as the outcome variable.

**Table S7. Differential abundance results for KEGG pathways level 2.** Model results were obtained by fitting LMMs on each pathway, while controlling for individual identity and unit membership. The estimate and p-value of the fixed effects are reported.

**Table S8. Differential abundance results for KEGG pathways level 3.** Model results were obtained by fitting LMMs on each pathway, controlling for individual identity and unit membership. The estimate and p-value of the fixed effects are reported.

**Table S9. Predictors of Shannon index, observed richness and Faith's phylogenetic diversity (PD) in females only.** Estimates with p-values <0.05 are highlighted in bold.

**Table S10. Predictors of the structure of the female gelada gut microbiome.** We carried out a PERMANOVA using 10,000 permutations and the Aitchison dissimilarity distance between samples.

**Table S11. Number of differentially abundant taxa for each predictor in the female samples.**

**Table S12. Differential abundance results for female samples at five taxonomic levels.**

**Table S13. Differential abundance results for KEGG pathways level 2 for female samples.** Model results were obtained by fitting LMMs on each pathway, while controlling for individual identity and unit membership. The estimate and p-value of the fixed effects are reported.

**Table S14. Differential abundance results for KEGG pathways level 2 for female samples.** Model results were obtained by fitting LMMs on each pathway, while controlling for individual identity and unit membership. The estimate and p-value of the fixed effects are reported.

**Table S15. Predictors of Shannon index, observed richness, and Faith's phylogenetic diversity (PD) on (1) all samples and (2) female samples on a rarefied dataset.** Parameters and tests are based on 758/439 samples and 131/70 individuals in all models. The LMMs were performed controlling for individual identity and unit membership. The 95% confidence intervals that do not cross zero and p-values of statistically significant results are highlighted in bold.

**Table S16. Results of PERMANOVA testing for the effects that significantly structure the gut microbiome of geladas on a rarefied dataset for (1) all samples or (2) female samples only. We used Bray Curtis, unweighted Unifrac or weighted Unifrac distances to assess between-sample dissimilarity.** 10000 permutations were carried out and individual identity was added as a strata in the model. The R-squared values indicate the amount of between-sample variation explained by each variable
